# The cortico-thalamic loop attunes competitive lateral interactions across retinotopic and orientation preference maps

**DOI:** 10.1101/2022.12.19.521069

**Authors:** Domenico G. Guarino, Andrew P. Davison, Yves Frégnac, Ján Antolík

**Affiliations:** Department for Integrative and Computational Neuroscience (ICN), Paris-Saclay Institute of Neuroscience (NeuroPSI); Campus CEA Saclay, 151 route de la Rotonde, 91400 Saclay, France; Computational System Neuroscience Group, Charles University, Malostranské nám. 25, 11800 Prague, Czechia

## Abstract

In the early visual system, corticothalamic feedback projections greatly outnumber thalamocortical feedforward projections. Extensive experimental and modeling work has been devoted to the functional impact of the feedforward pathway, but the role of its denser feedback counterpart remains elusive. Here, we propose a novel unifying framework where thalamic recurrent interactions and corticothalamic feedback act in a closed-loop fashion to attune multiple stimulus representations. At each position of the visual field, the loop puts into competition local representations of the stimulus in thalamus and cortex through direct excitation of narrow topologically-aligned portions of the thalamus, accompanied with peri-geniculate nucleus mediated broad inhibition suppressing the topological surround. We built a detailed conductance-based spiking model incorporating retinal input, lateral geniculate nucleus, peri-geniculate nucleus, primary visual cortex, and all the relevant intra-areal and feedback pathways. For the first time we perform comparative analyses between model configurations with completely or locally inactivated cortico-thalamic feedback, as in the experimental preparations. The model mechanistically explains (i) the existence of intra-thalamic surround suppression, (ii) the sensitivity of thalamic neurons to orientation tuning, (iii) the cortex-dependent center-surround opponency in thalamic cells, (iv) the cortical increase of size and orientation selectivity, (v) the cortically enhanced competition between cross-oriented domains within the hypercolumn, and (vi) the selective suppression of cortical functional connectivity. Our results integrate decades of experimental and theoretical research, supporting the hypothesis that cortico-thalamic loop exerts competitive influence between neighboring regions in the thalamus and cortex, complementing the lateral intra-V1 interactions in center-surround contextual modulation.

## Introduction

The classic framework to understand visual cortical function has relied dominantly on the feedforward thalamo-cortical drive (***Alonso et al., 1996***; ***Ferster and Miller, 2000***; ***Carandini et al., 2005***) and its modulation by local and distal cortico-cortical interactions (***Hubel and Wiesel, 1965***; ***Sillito et al., 1980***; ***Monier et al., 2008***; ***Fournier et al., 2014***). This view neglects the many functional observations that indicate significant participation of intrathalamic (***Usrey and Alitto, 2015***; ***Ghodrati et al., 2017***) and cortico-thalamic feedback influences in cortical computations (***Tsumoto and Suda, 1980***; ***Sherman and Koch, 1986***; ***Wörgötter et al., 1998***; ***Alitto and Usrey, 2008***; ***Briggs and Usrey, 2007***; ***Basso et al., 2005***). “*To adopt this attitude, however, indicates a failure to rise to the challenge of defining the critical questions to ask of the thalamus*” (***Jones, 1985***, p. 820).

Here we hypothesize that the recurrent interactions within the thalamus and the feedback loop between the cortex and thalamus are indispensable for understanding the mechanism through which stimulus representations are refined globally across the early visual system. These loops contribute to emerging properties in both thalamic and cortical responses and are instrumental to the interactions operating across the cortical retinotopic map. In particular, the cortico-thalamic loop can put into competition cortical encoding of stimuli at nearby retinotopic visual field locations through a ubiquitous connectivity motif of narrow excitation and broad inhibition. With direct excitation from V1 to LGN, cortical encoding of the visual stimulus at each retinotopic position projects its own attuned selectivity back to the thalamus (***Jones and Sillito, 1994***; ***Alonso et al., 1996***; ***Andolina et al., 2007***; ***Bijanzadeh et al., 2018***). With indirect - comparatively broader - inhibition mediated by the thalamic reticular nucleus, cortical stimulus encoding at each retinotopic position suppresses its topological surround (***Tsumoto et al., 1978***; ***Funke and Eysel, 1998***; ***Sillito and Jones, 2002***; ***Born et al., 2021***).

To test this hypothesis, we have built a detailed conductance-based spiking network model incorporating all the key elements of the higher-mammalian cortico-thalamic loop. Crucially, the model takes into account the peri-geniculate nucleus (PGN) - a major source of broad thalamic inhibition (***Jones, 1985***; ***Lam and Sherman, 2005***) - previously neglected in models of the cortico-thalamic loop (***Bonin et al., 2005***; ***Einevoll and Plesser, 2012***; ***Born et al., 2021***). In combination with the inclusion of experimentally established direct narrow cortico-thalamic excitation, we model the narrow-excitation/broader-inhibition motif of intra-thalamic and corticothalamic connectivity. To the thalamo-cortico model, we determine a minimal set of parameters that reproduces simultaneously a broad range of known functional properties of cortical and thalamic neurons under multiple stimulation conditions. With this set of parameters fixed, we then proceed to study how a range of experimental findings can be underpinned by the cortico-thalamic loop. For this purpose, we perform a comparative analysis across multiple spiking model configurations that mimic previously reported experimental inactivation conditions. We compare full model of the early visual system - an analog to intact brain condition - with two partially deafferented model configurations mimicking experimental preparations: (a) “local cortical feedback inactivation” configuration, where the cortical feedback was modulated by cortical current injection, and (b) “no cortical feedback” configuration, where the cortical feedback was completely absent.

This is the first model to reproduce a range of phenomena that previously lacked mechanistic explanation, including: the origin of extra-classical suppression in LGN independent of the cortex (***Cleland et al., 1983***; ***Alitto and Usrey, 2008***), the bias for stimulus orientation in LGN (***Daniels et al., 1977***; ***Creutzfeldt and Nothdurft, 1978***; ***Vidyasagar and Urbas, 1982***), the cortical enhancement of center-surround opponency in LGN (***Jones et al., 2012***). Our model also makes a range of testable predictions, including: (i) the role of the cortico thalamic loop in the enhancement of cortical selectivity for stimulus size and orientation, (ii) orientation-dependent and lateral competition within the hypercolumn, and (iii) stimulus size dependent modulation of center-surround effective interactions.

Overall this study demonstrates the importance of cortico-thalamic loop for shaping the evoked activity in V1. In contrast to the feed-forward and cortex-centric view currently dominating in neuroscience of vision, we show that a broad range of both thalamic and cortical functional properties can be substantially re-shaped by the cortico-thalamic loop, and hence we advocate for the necessity to rethink the encoding of stimulus in the early visual system in terms of a dynamic recurrent action concerted between the cortex and the thalamus.

## Results

We have built a conductance-based spiking model of the cat cortico-geniculate loop. The model was constrained by experimentally determined connectivity statistics, in-vitro and in-vivo single-cell electrophysiological properties, and functional measures of the evoked firing statistics in the intact and lesioned early visual system of the adult cat (***Figure 1***, see ***Table 1*** in the Methods section for the full list of model parameters and associated references).

**Table 1.**
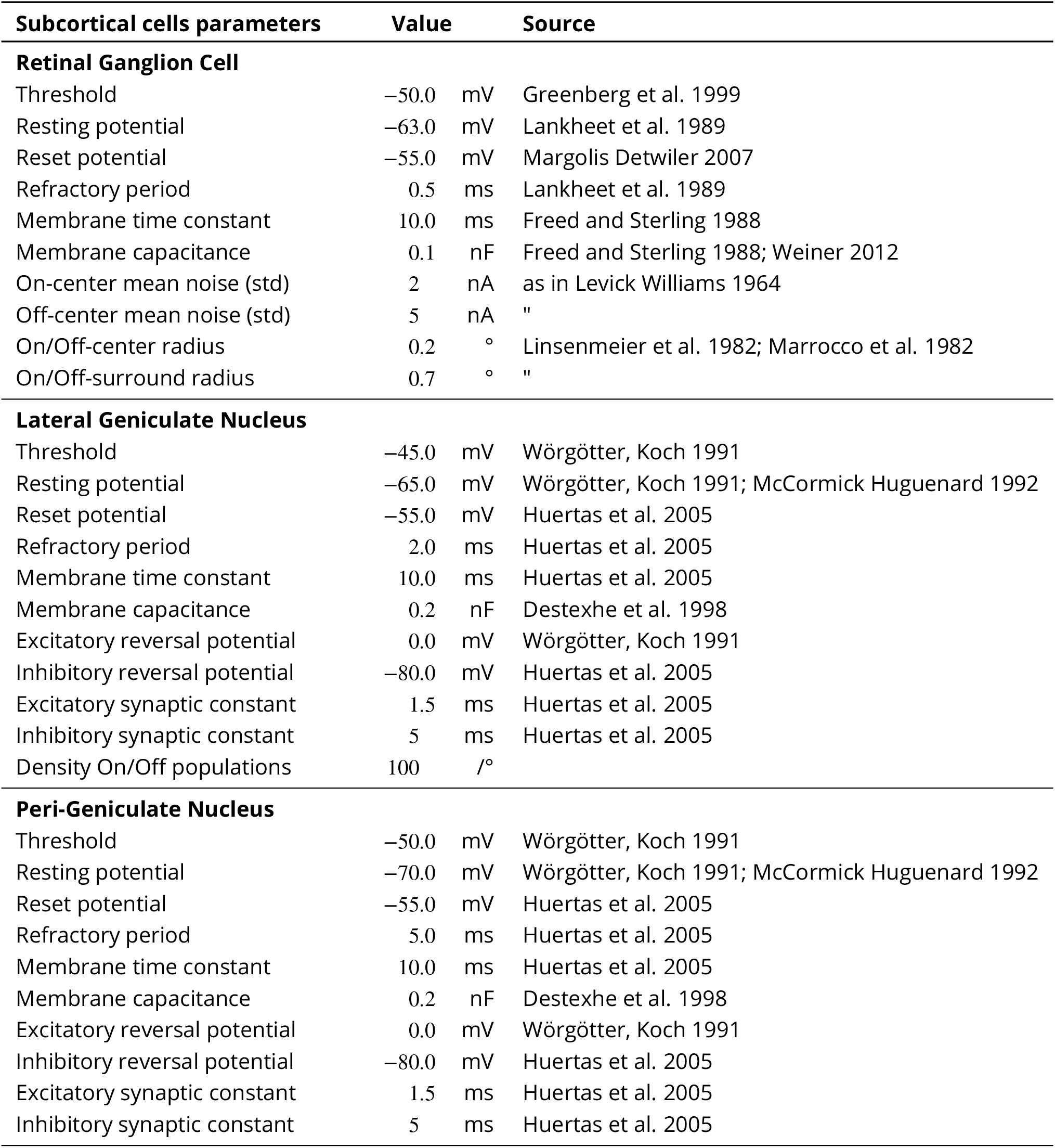
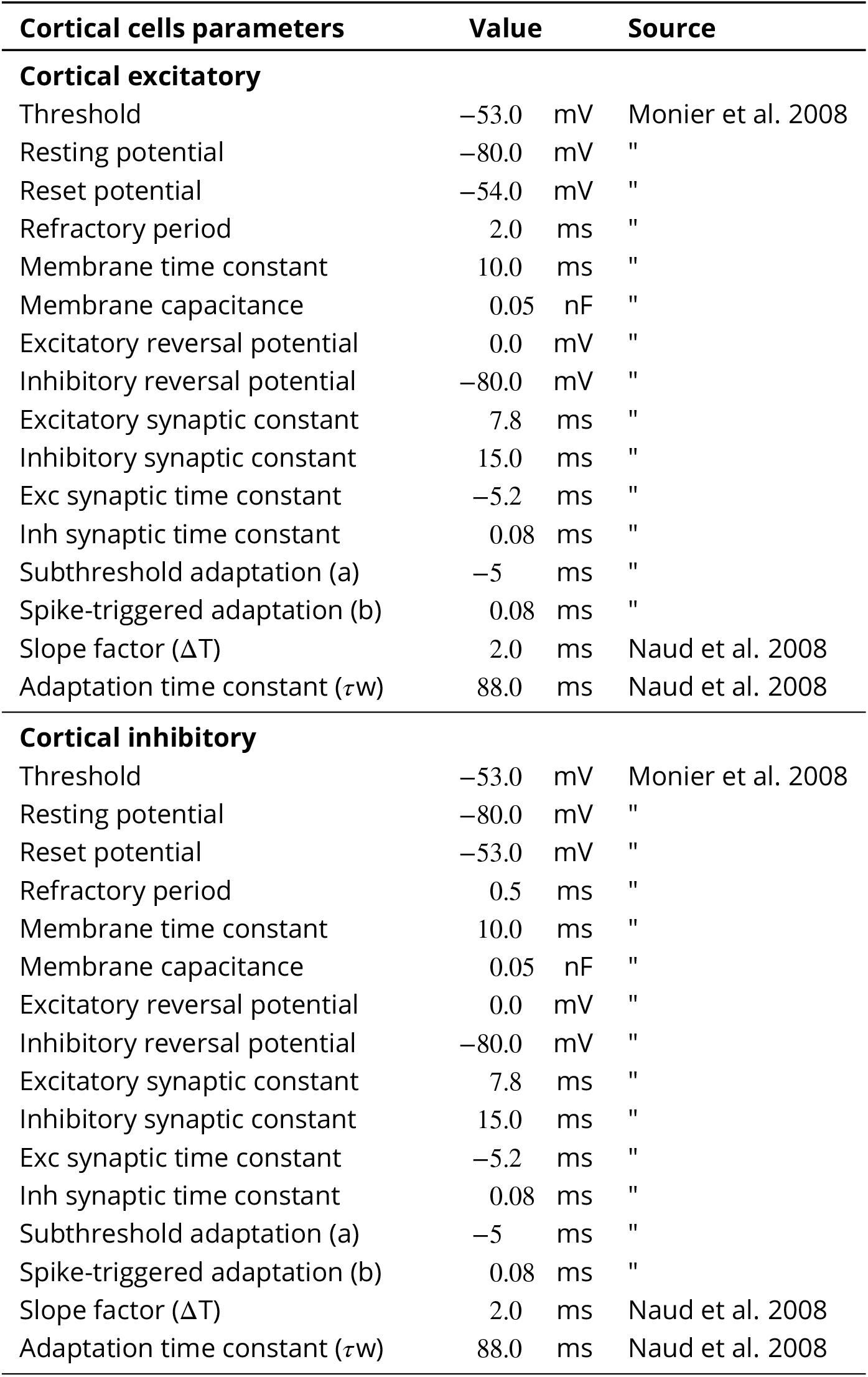
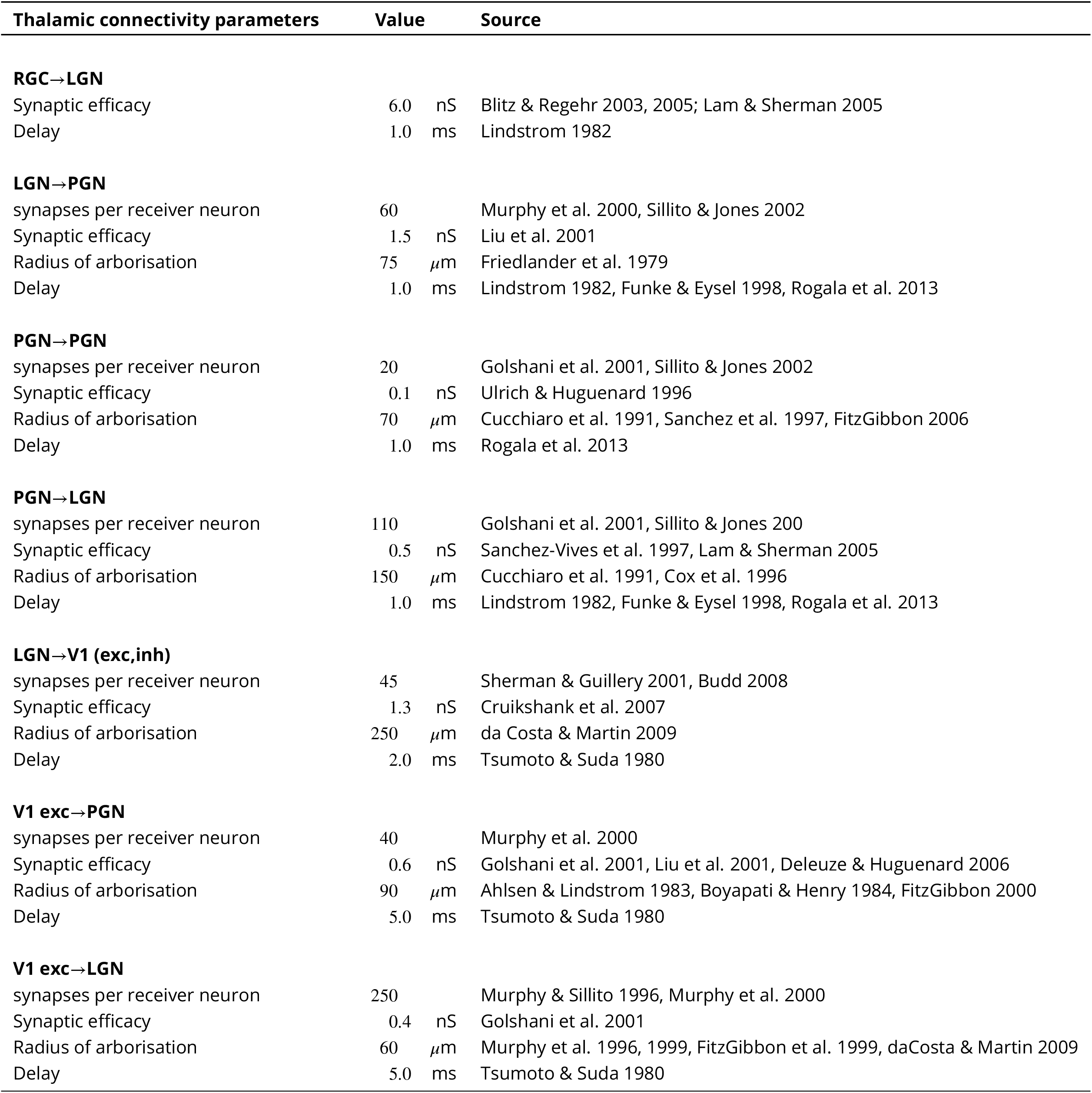
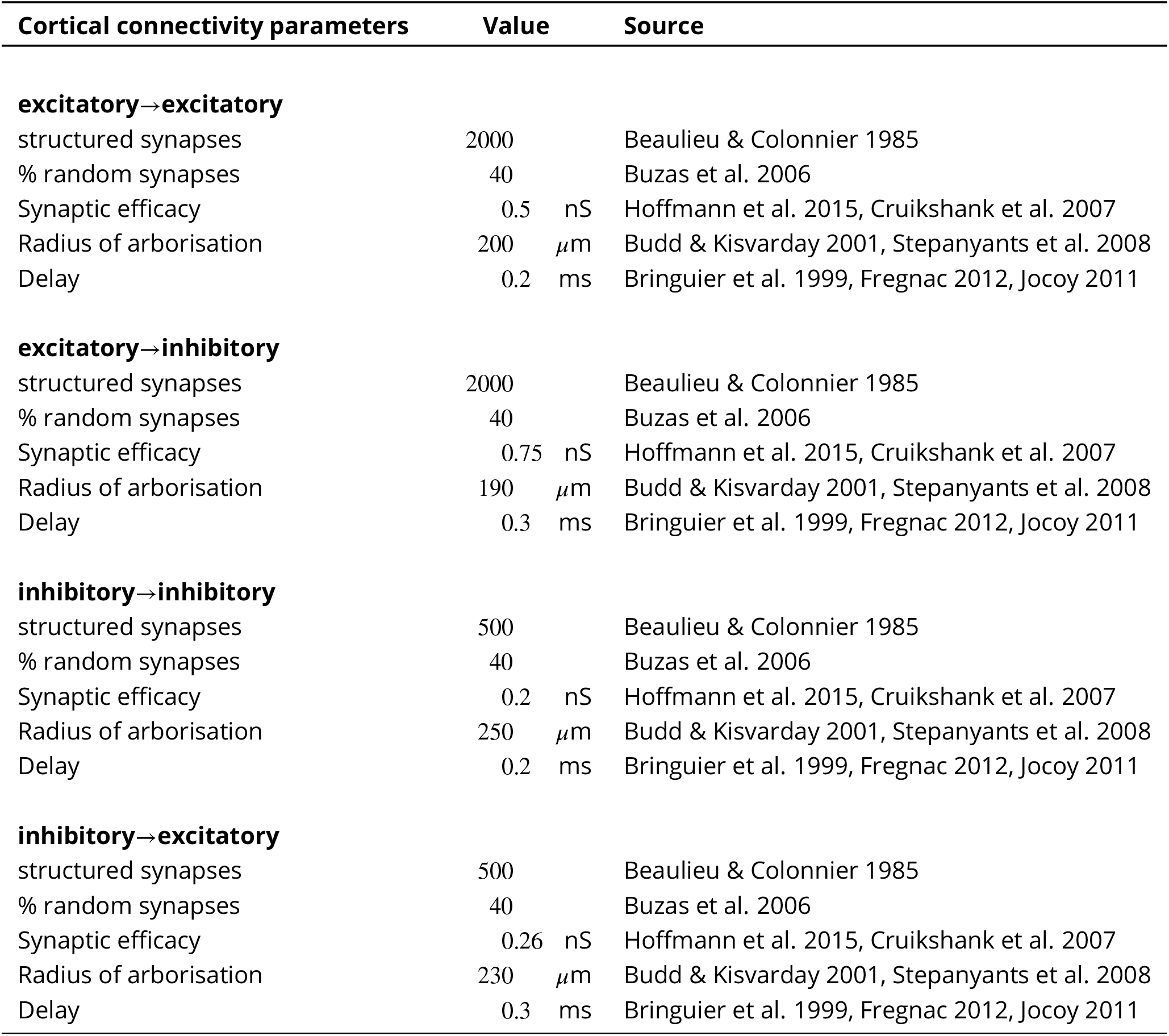
Model parameters.

| Subcortical cells parameters | Value |  | Source |
| --- | --- | --- | --- |
| **Retinal Ganglion Cell** |  |  |  |
| Threshold | −50.0 | mV | Greenberg et al. 1999 |
| Resting potential | −63.0 | mV | Lankheet et al. 1989 |
| Reset potential | −55.0 | mV | Margolis Detwiler 2007 |
| Refractory period | 0.5 | ms | Lankheet et al. 1989 |
| Membrane time constant | 10.0 | ms | Freed and Sterling 1988 |
| Membrane capacitance | 0.1 | nF | Freed and Sterling 1988; Weiner 2012 |
| On-center mean noise (std) | 2 | nA | as in Levick Williams 1964 |
| Off-center mean noise (std) | 5 | nA | " |
| On/Off-center radius | 0.2 | ° | Linsenmeier et al. 1982; Marrocco et al. 1982 |
| On/Off-surround radius | 0.7 | ° | " |
| **Lateral Geniculate Nucleus** |  |  |  |
| Threshold | −45.0 | mV | Wörgötter, Koch 1991 |
| Resting potential | −65.0 | mV | Wörgötter, Koch 1991; McCormick Huguenard 1992 |
| Reset potential | −55.0 | mV | Huertas et al. 2005 |
| Refractory period | 2.0 | ms | Huertas et al. 2005 |
| Membrane time constant | 10.0 | ms | Huertas et al. 2005 |
| Membrane capacitance | 0.2 | nF | Destexhe et al. 1998 |
| Excitatory reversal potential | 0.0 | mV | Wörgötter, Koch 1991 |
| Inhibitory reversal potential | −80.0 | mV | Huertas et al. 2005 |
| Excitatory synaptic constant | 1.5 | ms | Huertas et al. 2005 |
| Inhibitory synaptic constant | 5 | ms | Huertas et al. 2005 |
| Density On/Off populations | 100 | /° |  |
| **Peri-Geniculate Nucleus** |  |  |  |
| Threshold | −50.0 | mV | Wörgötter, Koch 1991 |
| Resting potential | −70.0 | mV | Wörgötter, Koch 1991; McCormick Huguenard 1992 |
| Reset potential | −55.0 | mV | Huertas et al. 2005 |
| Refractory period | 5.0 | ms | Huertas et al. 2005 |
| Membrane time constant | 10.0 | ms | Huertas et al. 2005 |
| Membrane capacitance | 0.2 | nF | Destexhe et al. 1998 |
| Excitatory reversal potential | 0.0 | mV | Wörgötter, Koch 1991 |
| Inhibitory reversal potential | −80.0 | mV | Huertas et al. 2005 |
| Excitatory synaptic constant | 1.5 | ms | Huertas et al. 2005 |
| Inhibitory synaptic constant | 5 | ms | Huertas et al. 2005 |

| Cortical cells parameters | Value |  | Source |
| --- | --- | --- | --- |
| **Cortical excitatory** |  |  |  |
| Threshold | −53.0 | mV | Monier et al. 2008 |
| Resting potential | −80.0 | mV | " |
| Reset potential | −54.0 | mV | " |
| Refractory period | 2.0 | ms | " |
| Membrane time constant | 10.0 | ms | " |
| Membrane capacitance | 0.05 | nF | " |
| Excitatory reversal potential | 0.0 | mV | " |
| Inhibitory reversal potential | −80.0 | mV | " |
| Excitatory synaptic constant | 7.8 | ms | " |
| Inhibitory synaptic constant | 15.0 | ms | " |
| Exc synaptic time constant | −5.2 | ms | " |
| Inh synaptic time constant | 0.08 | ms | " |
| Subthreshold adaptation (a) | −5 | ms | " |
| Spike-triggered adaptation (b) | 0.08 | ms | " |
| Slope factor ($\Delta T$) | 2.0 | ms | Naud et al. 2008 |
| Adaptation time constant ($\tau_w$) | 88.0 | ms | Naud et al. 2008 |
| **Cortical inhibitory** |  |  |  |
| Threshold | −53.0 | mV | Monier et al. 2008 |
| Resting potential | −80.0 | mV | " |
| Reset potential | −53.0 | mV | " |
| Refractory period | 0.5 | ms | " |
| Membrane time constant | 10.0 | ms | " |
| Membrane capacitance | 0.05 | nF | " |
| Excitatory reversal potential | 0.0 | mV | " |
| Inhibitory reversal potential | −80.0 | mV | " |
| Excitatory synaptic constant | 7.8 | ms | " |
| Inhibitory synaptic constant | 15.0 | ms | " |
| Exc synaptic time constant | −5.2 | ms | " |
| Inh synaptic time constant | 0.08 | ms | " |
| Subthreshold adaptation (a) | −5 | ms | " |
| Spike-triggered adaptation (b) | 0.08 | ms | " |
| Slope factor ($\Delta T$) | 2.0 | ms | Naud et al. 2008 |
| Adaptation time constant ($\tau_w$) | 88.0 | ms | Naud et al. 2008 |

| Thalamic connectivity parameters | Value | Source |
| --- | --- | --- |
| **RGC→LGN** |  |  |
| Synaptic efficacy | 6.0 nS | Blitz & Regehr 2003, 2005; Lam & Sherman 2005 |
| Delay | 1.0 ms | Lindstrom 1982 |
| **LGN→PGN** |  |  |
| synapses per receiver neuron | 60 | Murphy et al. 2000, Sillito & Jones 2002 |
| Synaptic efficacy | 1.5 nS | Liu et al. 2001 |
| Radius of arborisation | 75 $\mu\text{m}$ | Friedlander et al. 1979 |
| Delay | 1.0 ms | Lindstrom 1982, Funke & Eysel 1998, Rogala et al. 2013 |
| **PGN→PGN** |  |  |
| synapses per receiver neuron | 20 | Golshani et al. 2001, Sillito & Jones 2002 |
| Synaptic efficacy | 0.1 nS | Ulrich & Huguenard 1996 |
| Radius of arborisation | 70 $\mu\text{m}$ | Cucchiari et al. 1991, Sanchez et al. 1997, FitzGibbon 2006 |
| Delay | 1.0 ms | Rogala et al. 2013 |
| **PGN→LGN** |  |  |
| synapses per receiver neuron | 110 | Golshani et al. 2001, Sillito & Jones 200 |
| Synaptic efficacy | 0.5 nS | Sanchez-Vives et al. 1997, Lam & Sherman 2005 |
| Radius of arborisation | 150 $\mu\text{m}$ | Cucchiari et al. 1991, Cox et al. 1996 |
| Delay | 1.0 ms | Lindstrom 1982, Funke & Eysel 1998, Rogala et al. 2013 |
| **LGN→V1 (exc,inh)** |  |  |
| synapses per receiver neuron | 45 | Sherman & Guillery 2001, Budd 2008 |
| Synaptic efficacy | 1.3 nS | Cruikshank et al. 2007 |
| Radius of arborisation | 250 $\mu\text{m}$ | da Costa & Martin 2009 |
| Delay | 2.0 ms | Tsumoto & Suda 1980 |
| **V1 exc→PGN** |  |  |
| synapses per receiver neuron | 40 | Murphy et al. 2000 |
| Synaptic efficacy | 0.6 nS | Golshani et al. 2001, Liu et al. 2001, Deleuze & Huguenard 2006 |
| Radius of arborisation | 90 $\mu\text{m}$ | Ahlsten & Lindstrom 1983, Boyapati & Henry 1984, FitzGibbon 2000 |
| Delay | 5.0 ms | Tsumoto & Suda 1980 |
| **V1 exc→LGN** |  |  |
| synapses per receiver neuron | 250 | Murphy & Sillito 1996, Murphy et al. 2000 |
| Synaptic efficacy | 0.4 nS | Golshani et al. 2001 |
| Radius of arborisation | 60 $\mu\text{m}$ | Murphy et al. 1996, 1999, FitzGibbon et al. 1999, daCosta & Martin 2009 |
| Delay | 5.0 ms | Tsumoto & Suda 1980 |

| Cortical connectivity parameters | Value | Source |
| --- | --- | --- |
| **excitatory→excitatory** |  |  |
| structured synapses | 2000 | Beaulieu & Colonnier 1985 |
| % random synapses | 40 | Buzas et al. 2006 |
| Synaptic efficacy | 0.5 nS | Hoffmann et al. 2015, Cruikshank et al. 2007 |
| Radius of arborisation | 200 $\mu\text{m}$ | Budd & Kisvarday 2001, Stepanyants et al. 2008 |
| Delay | 0.2 ms | Bringuier et al. 1999, Fregnac 2012, Jocoy 2011 |
| **excitatory→inhibitory** |  |  |
| structured synapses | 2000 | Beaulieu & Colonnier 1985 |
| % random synapses | 40 | Buzas et al. 2006 |
| Synaptic efficacy | 0.75 nS | Hoffmann et al. 2015, Cruikshank et al. 2007 |
| Radius of arborisation | 190 $\mu\text{m}$ | Budd & Kisvarday 2001, Stepanyants et al. 2008 |
| Delay | 0.3 ms | Bringuier et al. 1999, Fregnac 2012, Jocoy 2011 |
| **inhibitory→inhibitory** |  |  |
| structured synapses | 500 | Beaulieu & Colonnier 1985 |
| % random synapses | 40 | Buzas et al. 2006 |
| Synaptic efficacy | 0.2 nS | Hoffmann et al. 2015, Cruikshank et al. 2007 |
| Radius of arborisation | 250 $\mu\text{m}$ | Budd & Kisvarday 2001, Stepanyants et al. 2008 |
| Delay | 0.2 ms | Bringuier et al. 1999, Fregnac 2012, Jocoy 2011 |
| **inhibitory→excitatory** |  |  |
| structured synapses | 500 | Beaulieu & Colonnier 1985 |
| % random synapses | 40 | Buzas et al. 2006 |
| Synaptic efficacy | 0.26 nS | Hoffmann et al. 2015, Cruikshank et al. 2007 |
| Radius of arborisation | 230 $\mu\text{m}$ | Budd & Kisvarday 2001, Stepanyants et al. 2008 |
| Delay | 0.3 ms | Bringuier et al. 1999, Fregnac 2012, Jocoy 2011 |

**Figure 1.**
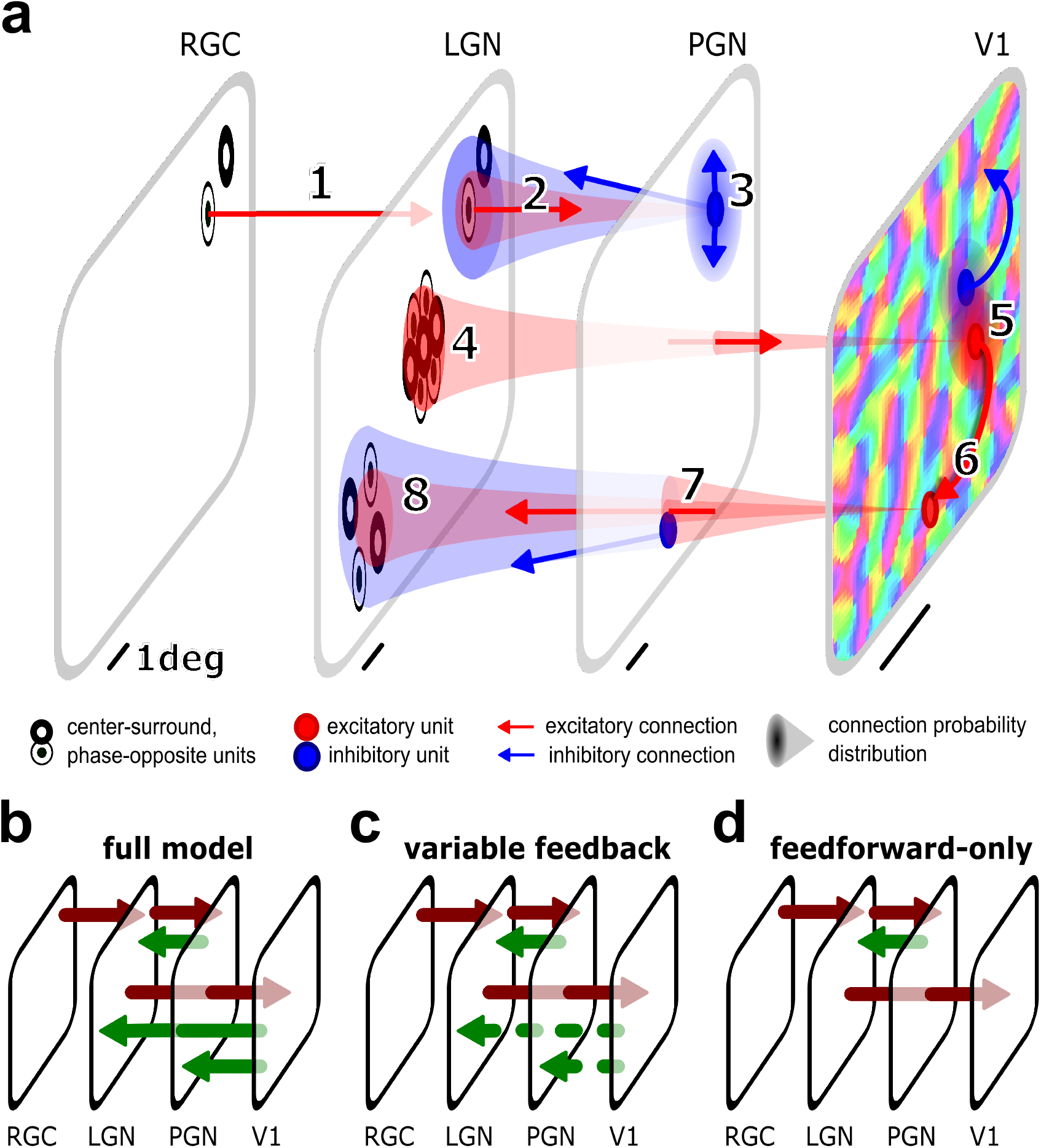
Model of the early visual system. (**a**) Model components with spatial and functional connectivity. From the left: Retina is modeled as a layer of phenomenological X retinal ganglion cells (RGC) of the two functional subgroups with center-surround opponency (ON-center, white inner disk, OFF-center, dark inner disk). Each RGC excites (red arrow **1**) one LGN cell determining its receptive field. Both ON- and OFF-center LGN cells excite (red arrow **2**) PGN cells (blue filled circles) and, in return, PGN cells inhibit (blue arrow **2**) both ON-and OFF-center LGN cells. PGN cells also inhibit each other (blue arrows **3**). In the primary visual cortex (V1), the afferent connections of excitatory (red circles) and inhibitory (blue circles) cells are formed by sampling connections from a Gabor probability distribution centered at the retinotopic position of the given cortical neuron and overlaid onto the sheets of ON- and OFF-center LGN cells (red cone and arrow **4**). Note that, in reality, LGN thalamocortical connections make collaterals into PGN, but for clarity we separated them into 2, 3, and 4. Positive Gabor subfields are overlaid on ON-center and negative on OFF-center sheets. V1 excitatory-to-inhibitory and inhibitory-to-excitatory connections (**5**) implement a push-pull connectivity (Troyer et al. 1998). The slow recruitment of lateral connections incorporates distance-dependent delays (**6**). Cortical feedback excitatory connections to the thalamus (red arrow) are formed by sampling connections from Gaussian probability distributions overlaid on PGN (**7**) and LGN (**8**) populations (see Methods). (**b-d**) Model configurations mimicking experimental preparations. (**b**) Full-model configuration, where all feedforward (purple) and feedback (green) pathways are present. (**c**) Variable cortical feedback configuration, where the cortical feedback was modulated by cortical current injection. (**d**) Feedforward-only configuration, where the cortical feedback was absent.

We tuned the model through an iterative workflow (see Methods) to match, qualitatively and quantitatively, the experimentally measured tuning of V1 firing rates to five features (contrast, spatial and temporal frequency, orientation, size) characterizing the test visual input (here, a full-field sinusoidal drifting grating), while restricting the free parameters within bounds outlined by existing experimental data. Through this iterative procedure, we found a single parameter set that satisfied all the available experimental constraints (***Figure 2***, and supplementary figures 1 and 2). Thenceforth, we no longer modified the optimized parameter set.

**Figure 2.**
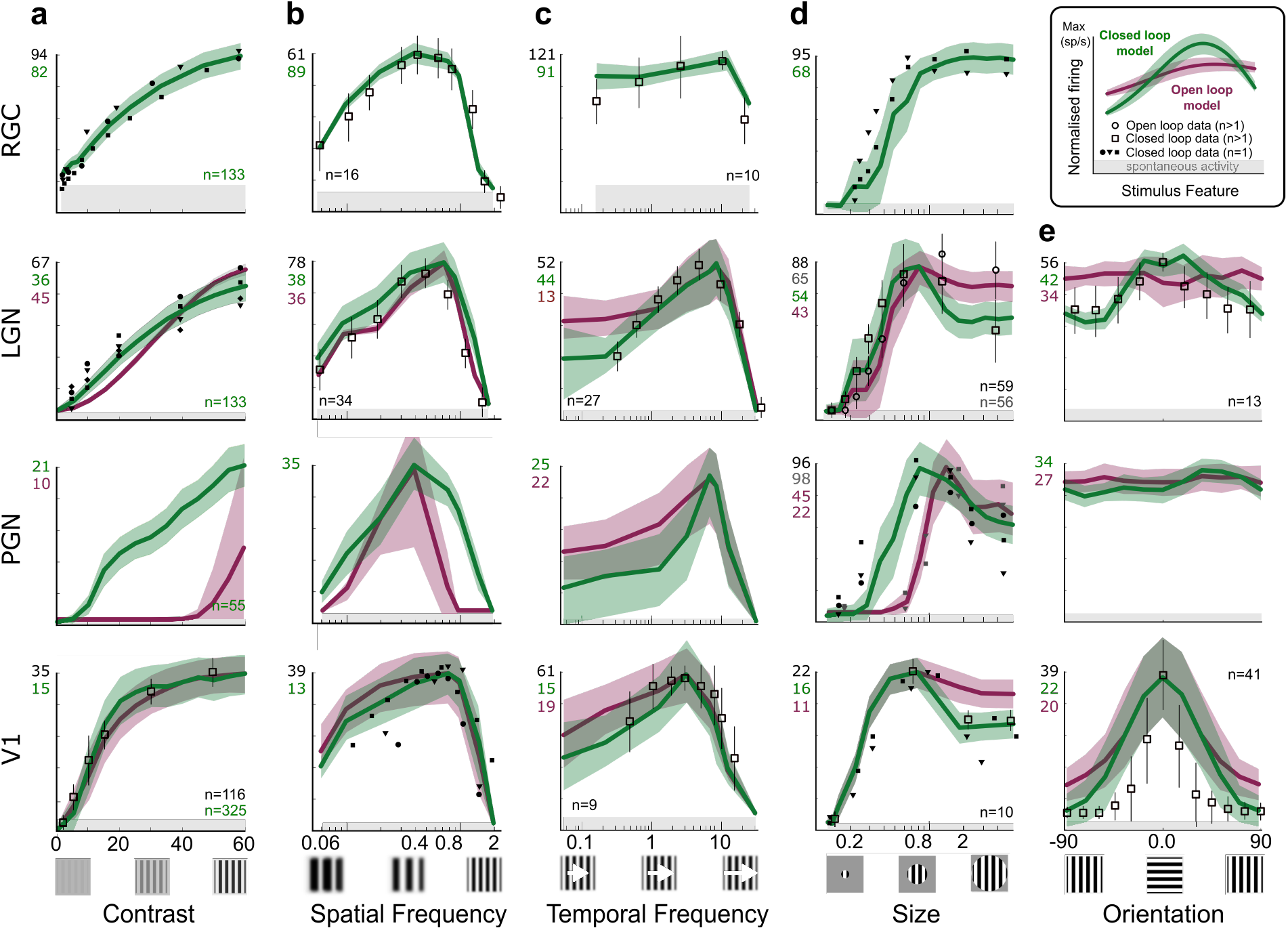
Experimental Data-driven constraints and optimized model fits. To tune our model, normalized trial-averaged firing rates from neurons virtually recorded in the closed (green curves) and open (purple curves) loop versions of our model were iteratively compared with closed and open loop experimental data, for five stimulus features variations: contrast (column **a**), spatial (**b**) and temporal (**c**) frequency, size (**d**), and orientation (**e**). Each row details a given integration stage in the early visual system (from top to bottom rows: RGC, LGN, PGN, V1). When available, experimental population averages (empty squares and circles with error bars) were used, otherwise, multiple single-cell recordings (various filled symbols) were used. The original firing rates are reported on the top of the vertical axis, using the same color code. The number of recorded model cells is reported in the lower right of column a (green), and, when available from experimental studies, in the other plots (black). When available, experimental data having multiple stimulus feature variations in the same study were used over data from single feature studies. We resorted to macaque data when data for cat was unavailable (see the stimuli section of Methods for experimental data sources).

We then proceeded to dissect the functional impact of corticothalamic feedback by comparing cortical and thalamic responses to the different sets of stimuli, between the full closed-loop configuration (Figure 1b), against the open loop and altered feedback configurations (Figure 1c, d). These model configurations correspond to the majority of experimental preparations utilized in studies of the cortico-thalamic loop. In the following, we organize the results around the classical view of the two integration loci for the cortico-thalamic-loop: the thalamic and cortical viewpoints. However, we will remind the reader of the full integration of this system by making punctual links between figures from one viewpoint to the other.

### The Thalamic viewpoint

Two debated questions in the experimental literature – (a) the origin of LGN extra-classical surround suppression, and (b) the origin of LGN orientation bias – offer the opportunity to distinguish the functional role of corticothalamic feedback from thalamic selectivity supported by the input from the retina.

#### Cortical feedback enhances thalamic selectivity for stimulus size

In the early visual pathway, neural responses generally vary with the size of a contrast-defined stimulus. For small sizes, the response increases as excitation is dominant. For stimuli extending beyond the classical receptive field, responses saturate and then decrease because of gradual recruitment of surround inhibition. Such surround suppression has been also observed in LGN neurons (***Cleland et al., 1983***). The origin of this effect is still a matter of debate. When the primary visual cortex feedback is disabled, the surround suppression in LGN principal cells declines in comparison to the intact condition but is not fully eliminated (***Murphy and Sillito, 1987***; ***Jones et al., 2000***; ***De Labra et al., 2007***; ***Alitto and Usrey, 2008***; ***Usrey and Alitto, 2015***). In our model, we found that the corticothalamic feedback enhances an already existing selectivity for stimulus sizes that originates in the narrow projections from the LGN to its surrounding peri-geniculate nucleus (PGN), reciprocated by broader inhibitory projections from PGN to LGN.

Specifically, in our virtual experiments, we stimulated the different model configurations with circular patches of drifting gratings of varying radii (see Methods), and we recorded membrane potentials, spikes, and input conductances of virtual LGN and PGN cells. In the absence of any cortical feedback, the modeled thalamus already exhibited extra-classical surround suppression of LGN responses (***Figure 3***a). The full model exhibited a significant increase in mean response at small sizes (+16.1%, Welch t-test, p<0.001), with a significant decrease in mean response for large sizes (−18.3%, Welch t-test, p<0.001). This results in a significant reduction of the average suppression index between full and feedforward-only configurations (−43.3%, from 0.29±0.01 to 0.16±0.01, Welch t-test, p<0.001).

**Figure 3.**
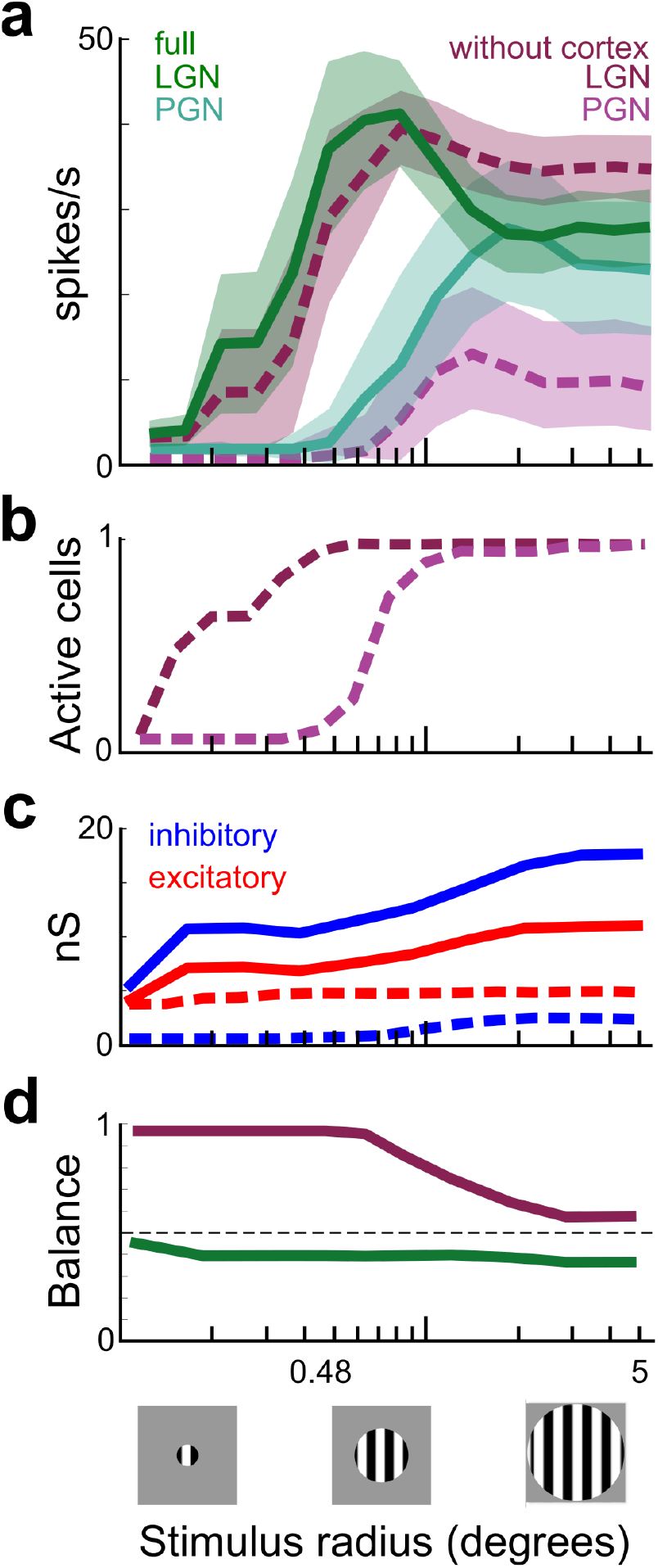
Cortical feedback increases thalamic response selectivity for sizes. (**a**) Population trial-averaged size tuning curves (shaded SEM) for virtual neurons, recorded in LGN (n=243, green) and PGN (n=57, purple), respectively in the feedforward-only model configuration (light dashed lines and shades), and in the full model configuration (dark solid lines and shades). Note, in the full model, the shift in the preferred stimulus size for LGN RFs, together with a significantly lower plateau. (**b**) Normalized count of feedforward-only LGN and PGN active cells as a function of stimulus size. PGN cells require larger sizes to be recruited. (**c**) Trial-averaged size tuning of mean excitatory (red) and inhibitory (blue) conductances in the LGN. In the feedforward-only model configuration (dashed), the LGN was dominated by excitatory conductance. The inhibitory conductance increased beyond 0.48° stimulus radius, corresponding to the PGN increase in excitatory inputs from the LGN. In the full model configuration (solid), the LGN was dominated by inhibitory conductance. (**d**) Excitatory-to-inhibitory balance in the LGN. In the feedforward-only model configuration (dashed), the LGN conductance ratio is dominated by excitation and becomes balanced only at large sizes. In the full model configuration (solid), it is overall more balanced and dominated by inhibition.

The presence of surround suppression even in the absence of cortical feedback exhibited by the feedforward-only model configuration, and its reduction of suppression index when compared to the full model, are consistent with in-vivo experimental studies. ***Murphy and Sillito (1987***) recorded LGN X-type cells from cats with either intact or ablated primary cortical areas and found a significant reduction (−35.5%) of the average suppression index between control and ablated cortex conditions (from 0.71±0.02 to 0.43±0.02, t-test, p<0.001). ***Andolina et al. (2013***) reported a significant reduction of mean LGN responses for drifting grating stimuli of varying size (−20.9%, t-test p=0.017) when applying the GABA promoter muscimol over cat visual cortical areas. For the same type of stimulus, but without manipulation of the cortex, ***Usrey and Alitto (2015***) found a suppression index of 0.43±0.02 in LGN cells.

We next proceeded to dissect the underlying mechanisms of extra-classical surround suppression. In the feedforward-only model configuration, the percentage of LGN and PGN cells progressively recruited with the increase of stimulus size showed two different activation thresholds (***Figure 3***b). For small stimuli (below 0.5 deg), the percentage of LGN cells firing above ongoing activity followed the stimulus size, the number of recruited LGN cells remained below the spatial summation threshold of PGN cells, and, consequently, no retroactive inhibition from PGN was induced. In contrast, large stimuli (above 0.5 deg) recruited enough LGN cells to surpass the PGN activation threshold, and thus PGN began providing recurrent inhibition. The growth rate of PGN activity with respect to stimulus size was steeper (11.1 growth rate) than that of LGN. This means that for sufficiently large stimuli, PGN inhibition overtakes the additional stimulus size induced excitation in LGN cells, augmenting the strength of size-dependent suppression.

To get a mechanistic understanding of these results, we used our model to extract population-averaged input conductance tuning curves of LGN (***Figure 3***c). In the feedforward-only configuration, the excitatory conductance grew with the stimulus size, while inhibitory conductance started to grow only with larger stimuli, but then increased much faster following the increase in PGN firing. In contrast, in the full model configuration, inhibition was dominant and both excitation and inhibition increased concomitantly with stimulus size. To highlight the functional impact of the open vs. closed loop configuration on the size tuning of LGN neurons, in Figure 3d we analyzed the excitatory-to-inhibitory conductance balance (EICB) — the ratio of excitatory to total input conductances 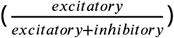, where values close to 0 represent inhibition-dominated regimes and values of 1 excitatory-dominated regimes. We showed that, in the feedforward-only model configuration, LGN cells are characterized by an excitatory-dominated regime for small sizes (EICB=0.91±0.08), which shifted towards inhibition at large sizes (EICB=0.62±0.12). This confirms that the inhibitory contribution from PGN required large-sized stimuli to be effective at inhibiting LGN response. In the full model, the corticothalamic feedback raises PGN inhibition, leading to a further shift toward inhibition (small: EICB=0.45±0.05, large: EICB=0.38±0.11).

Cortical studies have shown that both excitatory and inhibitory conductances are size tuned in V1 neurons, supporting the view that inhibition-stabilization is the underlying mechanism for such conductance tuning (***Ozeki et al., 2009***). In contrast, in LGN, our simulations predict that this is not the case. Instead, both excitatory and inhibitory conductances increase with stimulus size, while the size-tuning of the spiking response is mediated through stimulus-size-dependent changes to excitatory vs. inhibitory balance. Our model thus predicts a different mechanism of size tuning generation in LGN in comparison to V1.

These results suggest that intra-thalamic connectivity is sufficient per se to foster competition between spatially offset LGN neurons through the ubiquitous mechanism of short-range excitation and longer-range inhibition. Cortical feedback further enhances this competition. However, the cortex is also sensitive to other stimulus features, such as orientation, which may be projected onto the thalamic circuitry, as we will see in the following section.

#### Cortical feedback introduces a thalamic bias for stimulus orientation

Although orientation selectivity is considered a hallmark of thalamocortical convergence, weak sensitivity of cell responses to stimulus orientation has been reported for cat LGN cells (***Daniels et al., 1977***; ***Creutzfeldt and Nothdurft, 1978***; ***Vidyasagar and Urbas, 1982***; ***Vidyasagar et al., 2015***). Cortical feedback has been hypothesized to influence both the tuning and the distribution of orientation preference (***Vidyasagar and Urbas, 1982***; ***Krug et al., 2001***; ***Sedigh-Sarvestani et al., 2017***), but the mechanism of this latter effect remains unclear. In our model, we found that thalamic orientation bias is only expressed when the distribution of orientation preferences of cortical afferents to the thalamus is homogeneous. On the contrary, when the cortical feedback orientation preference is heterogeneous, no orientation preference is expressed in the thalamus.

Simulations were run, using full-field drifting grating stimuli of varying orientations, and recording membrane potentials, spikes, and input conductances from virtual LGN cells, in the feedforward-only and full model configurations. The predictions of our model agree with the available experimental data based on trial-averaged LGN firing rate responses. Similar effects of cortical deafferentation are found for the orientation bias and the ratio of preferred to non-preferred orientations (***Figure 4***a). Mean orientation bias significantly decreased (−15.7%, paired t-test, p=0.0007) from 1.27±0.14, in the feedforward-only configuration, compared to 1.13±0.07, in the “full” model. These results are in line with ***Vidyasagar and Urbas (1982***), who tested orientation selectivity in cat LGN cells, but using moving bars, and reported that biases of LGN X-cells changed significantly between the two configurations, with a 14.5% decrease of mean orientation bias from 1.83±0.55 to 1.74±0.62 (paired t-test, p<0.001).

**Figure 4.**
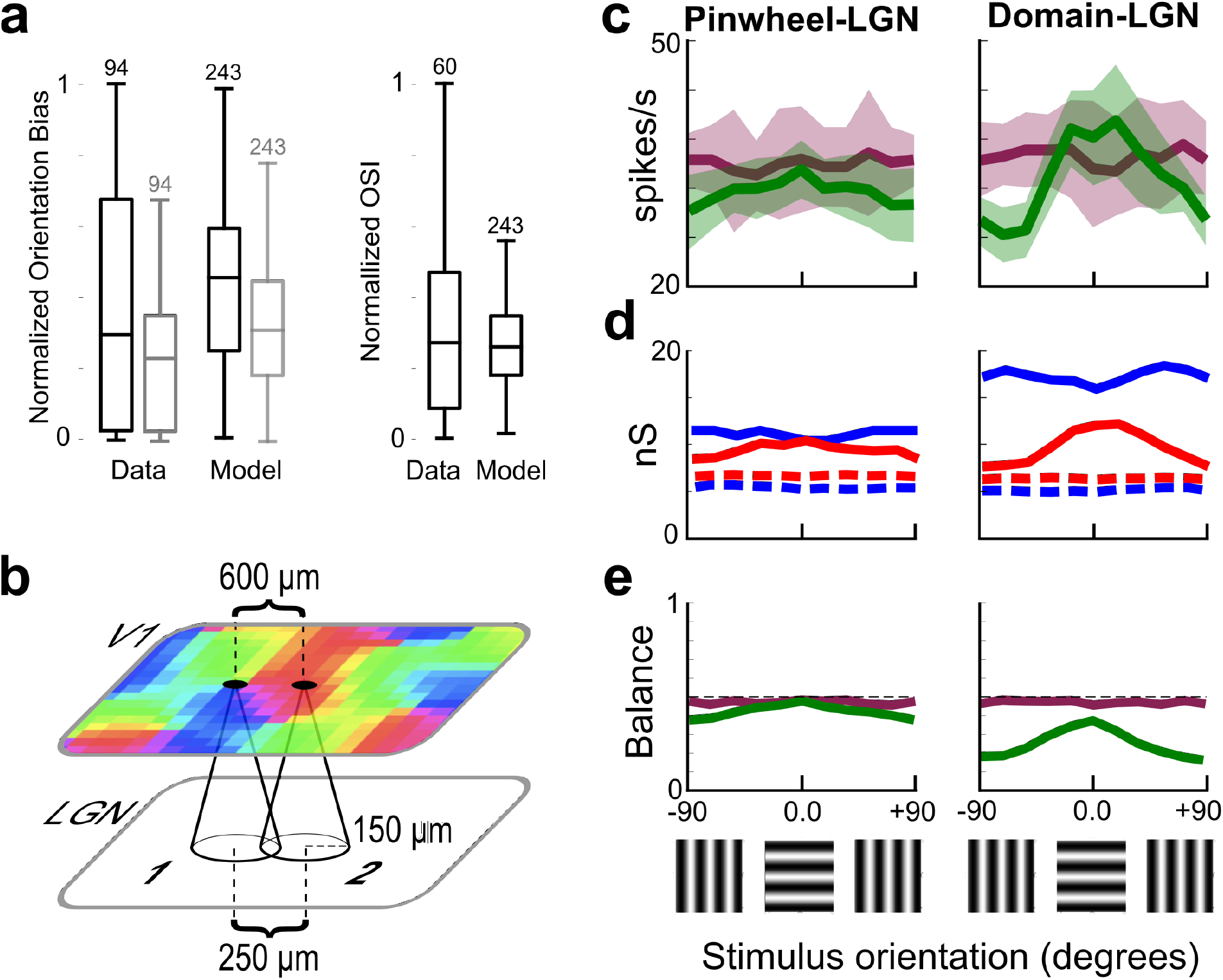
Cortical feedback introduces thalamic selectivity for orientation. (**a**) Left: Normalized orientation bias ratio in the full (black) and feedforward-only (gray) configurations, for (n=243) LGN cells. Left: Using drifting bars, experiments by Vidyasagar and Urbas (1982) showed a significant mean reduction of orientation bias ratio (n=94, drifting bars of 15×0.1 deg, moving at 5 deg/s). Our model also exhibited such reduction when measured using drifting gratings. Right: Orientation selectivity indexes measured using drifting gratings by Naito et al. 2007 (only available for intact condition). Our full model matches the OSI measured in cats when measured using an identical stimulus. (**b**) Setup of the virtual orientation tuning experiment. Two groups of model LGN cells were selected based on the orientation preference of their cortical feedback afferent inputs (1: from a cortical pinwheel, and 2: from a “0 degree” cortical ISO-domain). (**c**) For the LGN group receiving input exclusively from cortical pinwheels, the trial-averaged firing orientation tuning curves (top) show no significant selectivity changes between the full (green, shaded SEM) and “feedforward-only” model (purple) configurations. For the LGN group receiving selective inputs from 0 degrees-oriented ISO-preference cortical domains, the trial-averaged firing orientation tuning curves (top) showed selectivity in the full model (green), significantly different from the nonselective units recorded in the feedforward-only model configuration (purple). (**d**) The trial-averaged synaptic conductance tuning curves for both group 1 and 2 (dashed) confirm the absence of selectivity in the feedforward-only model configuration. The conductance tuning curves of the full model show tuned conductances for both groups. (**e**) Excitatory-to-inhibitory balance. In the feedforward-only model configuration (dashed), the LGN conductance input ratio is balanced for both groups. In the full model configuration (green), only for the Domain-LGN group, global input conductance is inhibitory dominated, becoming more balanced only when integrating the orientation-biased cortical input.

The orientation tuning of cat LGN cells in the presence of intact V1 has been tested also with sinusoidal drifting gratings (***Naito et al., 2007***; ***Suematsu et al., 2013***; ***Osaki et al., 2018***). All studies reported mean orientation selectivity indexes (OSI) — the normalized circular distance between stimuli and responses — for LGN X cells of 0.3 (***Suematsu et al., 2013***), in line with the OSI measured in the intact model (***Figure 4***a, right).

Guided by the hypothesis that the local cortical orientation preference (***Payne and Peters, 2002***) could differently bias thalamic responses, we divided the virtual LGN recordings into two groups (***Figure 4***b) such that, in the corresponding cortical location, the majority of cortical cells belonged to the same ISO-orientation preference domain (ISO-Domain-LGN group, where the cortical ISO-domain had 0 degrees preference), or expressed a mixture of preferences (Pinwheel-LGN group). In the feedforward-only model configuration, there was no significant response preference toward any orientation in either LGN groups (***Figure 4***c, blue lines). However, in the “full” model (green lines), the two groups of thalamic cells responded differently to the orientation protocol. We found statistically marginal change due to cortical feedback in the Pinwheel-LGN group (Welsh t-test, p=0.097). In contrast, a highly significant change in orientation preference emerged (Welsh t-test, p=0.0071) in the ISO-Domain-LGN group (***Figure 4***c).

A mechanistic explanation can be provided by comparing the excitatory and inhibitory conductance tuning curves of the two LGN cell groups in the feedforward-only and full model configurations (***Figure 4***d). The corticothalamic feedback significantly raised (Welsh t-test, p=0.0001) the excitatory conductance of the ISO-domain-LGN group, preferentially at the orientation matching that of the co-registered cortical domain (by convention, 0 degrees). It also raised the inhibitory conductance preferentially for cross-oriented stimuli. Similar biases, but of much smaller magnitude, were present in the conductances recorded from the Pinwheel-LGN group. We hypothesize that this residual orientation selectivity in the Pinwheel-LGN group is due to remnant non-uniformity in the orientation preferences represented around a pinwheel. The excitatory-to-inhibitory conductance balance (***Figure 4***e) recapitulates these orientation tuning characteristics in the ISO-domains vs. Pinwheel groups. In the feedforward-only model configuration, both LGN groups were characterized by a lack of impact of stimulus orientation on the E/I balance (Pinwheel-LGN: 0.48±0.1, Domain-LGN: 0.46±0.16; ***Figure 4***d,e). In the full model, the corticothalamic feedback selectively raised inhibition only in the ISO-Domain-LGN cell group, and more so for stimulus orientations orthogonal to the orientation preference of the retinotopically co-registered cortical domain (EICB Pinwheel-LGN: 0.38±0.15, Domain-LGN: 0.21±0.1; ***Figure 4***d,e). Overall these results demonstrate that the cortex can imprint orientation preference onto LGN neurons through cortico-thalamic feedback, but this phenomenon is dependent on the functional organization of the surrounding area of retinotopically co-registered cortex.

#### Cortical feedback enhances center-surround opponency in LGN cells

The experimental studies and our modeling presented in the previous two sections hint at the possibility that the functional influence of the corticofugal pathway onto its thalamic target could depend on the spatial relationship between thalamic and cortical receptive fields (RFs). In fact, the seminal work by ***Tsumoto et al. (1978***) showed that spatial overlap between the RFs of V1 cells and their target LGN cells determined the sign of the corticothalamic modulatory effect on LGN responses. Later, ***Jones et al. (2012***) recorded LGN cell responses to patches of drifting sinusoidal gratings of varying diameters, while reversibly inactivating local regions of V1, whose RFs were either overlapping or non-overlapping with those of the recorded LGN cells. They reported enhanced facilitation for overlapping LGN-V1 RFs, together with enhanced suppression for non-overlapping LGN-V1 RFs. However, the mechanism underlying such opponency remains unclear.

The constraints outlined by anatomical and functional studies of both intra-thalamic and corticothalamic connectivity, which we introduced in our model, imply the spatial impact of the disynaptic inhibitory pathway combining V1→PGN→LGN was broader than the direct monosynaptic excitation of V1→LGN connections. We, therefore, hypothesized that a stimulus that engages over-lapping cortical and thalamic receptive fields will induce a corticothalamic enhancement of the LGN center response through the direct excitatory pathway, as well as a concomitant enhancement of the suppressive surround through the broader indirect inhibitory pathway. We tested this hypothesis by replicating the experiments of ***Jones et al. (2012***) in our simulations and found that indeed the corticothalamic feedback enhances the effects of intra-thalamic center-surround opponency, demonstrating that the specific anatomical parameters of the V1→PGN→LGN can explain experimental data of ***Jones et al. (2012***).

We stimulated the model with circular patches of drifting gratings of varying radius and we inactivated spatially registered cortical excitatory cells (***Figure 5***a, b; see “variable-cortical-feedback” configuration, ***Figure 1***c). Pairs of cortical and thalamic neurons were grouped according to the distance between their RFs center in two categories: overlapping — when cortical RF subtended an area of visual space containing the center of the recorded LGN cell (***Figure 5***a), and non-overlapping — when the cortical RF subtended area of visual space near but offset from that sampled by the recorded LGN cell (***Figure 5***b).

**Figure 5.**
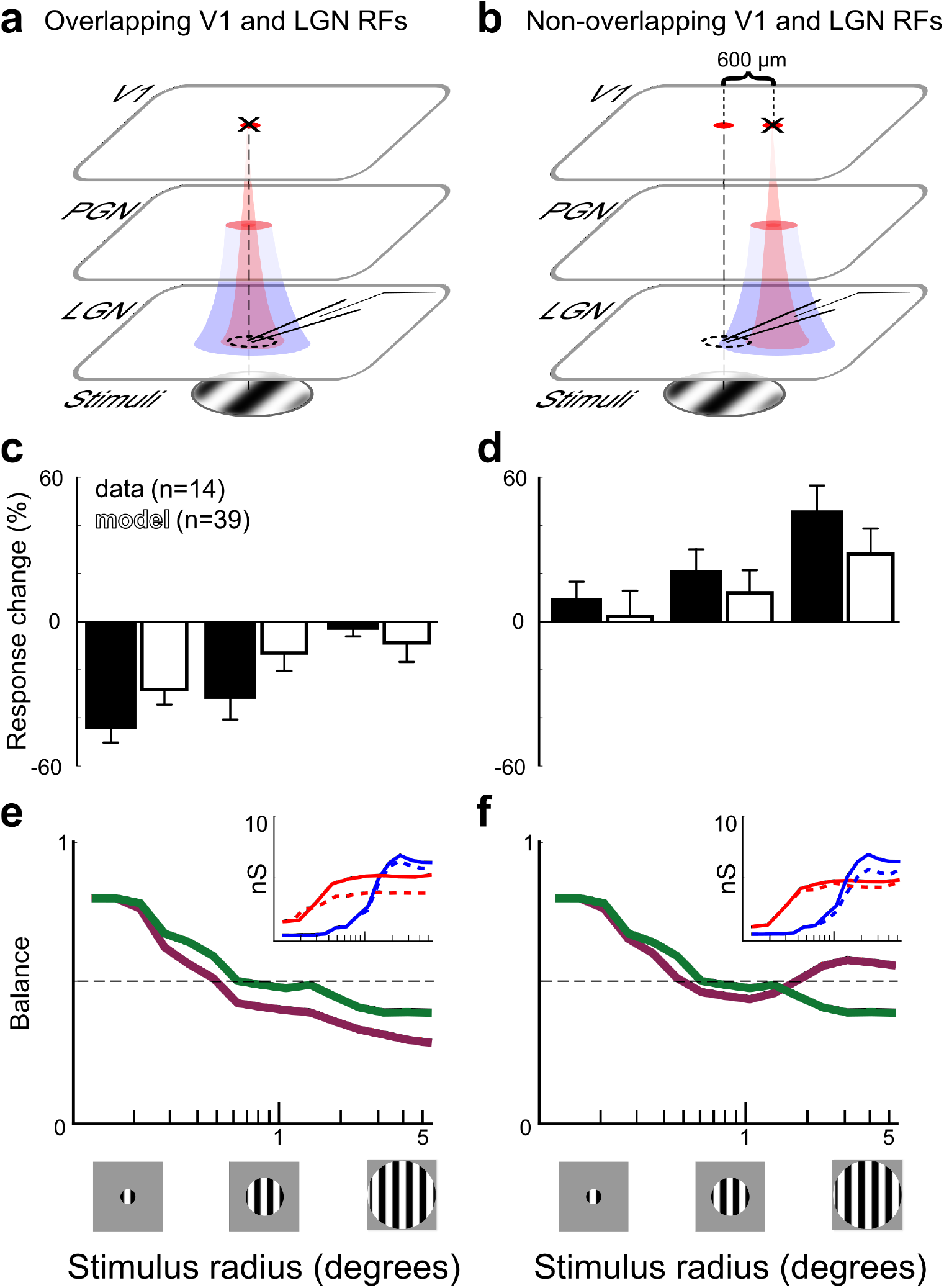
Cortical feedback enhances thalamic center-surround opponency. (**a,b**) Setup of the virtual experiment to study the dependence of size tuned LGN responses on the degree of retinotopic overlap with cortical receptive fields. As in Jones et al. 2012, cortical locations (red circles, ∼600 *μ*m apart) were chosen such that in one condition they were retinotopically overlapping (**a**, dashed black line) the RFs of recorded LGN cells (dashed circle), while in the second condition they were non-overlapping with (**b**, still in proximity to) the recorded LGN RFs. The distance between overlapping and non-overlapping locations was chosen such that the effective corticothalamic direct excitation (red cone) was flanked by a disynaptic indirect inhibition (blue annulus). The overlapping cortical location was inactivated in **a, c, e**, while the non-overlapping cortical location was inactivated in **b, d, f** (black cross over the corresponding red circle). (**c,d**) Population summary histogram of the mean percentage change in LGN cell responses between intact and locally-inactivated model cortex (empty bars), vs. Jones et al. 2012 data (filled bars). The signs of the changes in the model were opposite for the overlapping and non-overlapping conditions (negative in c and positive in d) and their magnitude decreased (**c**) or increased (**d**) with the grating stimulus radius (abscissa in degrees), in agreement with the experimental data. (**e**) Excitatory-to-Inhibitory conductance balance (EICB) is reduced for the overlapping inactivation (full model in green, local cortical inactivation in purple). (Inset) Trial-averaged mean excitatory (red) and inhibitory (blue) synaptic conductance tuning curves of model LGN cells. When the overlapping cortical location is inactivated, only the excitatory conductance is reduced appreciably (dashed), whereas no or minimal change is observed in the inhibitory conductance. (**f**) The EICB is increased only at large sizes during non-overlapping inactivation. (Inset) When a non-overlapping cortical site is inactivated, the inhibitory conductance is lower (dashed) while the excitatory conductance profile is unchanged.

Preferred stimulus size was defined as the grating patch size eliciting maximal response in the given LGN cell (peak of the size tuning-curve) recorded in the intact (control) condition. We then measured the percentage change of the peak response of LGN neurons (n=45) during cortical inactivation. The response changes were averaged for three grating patch size ranges: (i) smaller-than-preferred stimulus sizes, (ii) preferred stimulus size, and (iii) larger-than-preferred stimulus sizes, When we inactivated the overlapping cortical group (***Figure 5***c), the recorded LGN responses showed a significant decrease for less-than-preferred sizes (−29.3%, empty bars; Wilcoxon pair test, p=0.001), together with smaller decreases for preferred (−16.7%) and for larger-than-preferred sizes (−9.1%). In contrast, inactivation of the non-overlapping cortical group lead to a significant increase in response (+26.5%) for larger-than-preferred sizes (***Figure 5***d; Wilcoxon matched pair test, p=0.002), as well as smaller but positive changes for preferred (+13.2%) and smaller-than-preferred sizes (+2.4%).

Model results are in qualitative agreement with the reference experimental study by ***Jones et al. (2012***). In their work, the local inactivation of cortical cells having RF centers in overlap with recorded LGN RF centers resulted in a significant decrease of responses in those LGN neurons (−45.8%, Figure 5c, filled bar) below control levels for less-than-preferred sizes (Wilcoxon pair test, p=0.003). We found a smaller but statistically significant decrease for preferred and a statistically non-significant decrease for larger-than-preferred sized stimuli. In the work by ***Jones et al. (2012***), local inactivation of cortical cells having RFs non-overlapping with recorded LGN ones resulted in a significant increase (+46.3%, ***Figure 5***d, filled bars) above control levels in larger-than-preferred sizes (Wilcoxon pair test, p=0.003), together with a smaller increase for preferred and for smaller-than-preferred sizes. The smaller magnitude of changes due to cortical inactivation observed in the model relative to ***Jones et al. (2012***) might indicate that the feedback connection density and/or synaptic weights, both poorly constrained by existing literature, are under-estimated in the model.

We could dissect our model to gain a mechanistic understanding of the experimental results for the overlapping and non-overlapping configurations. We hypothesized that the connectivity constraints of our model would result in the cortical enhancements of thalamic center-surround opponency. This hypothesis was supported by the size-tuning curves of the excitatory and inhibitory input conductances. Our simulations showed indeed that, when suppressing cortical activation in regions in register with the activated thalamic receptive fields (***Figure 5***e), the withdrawal of cortical feedback excitation led to a reduced thalamic response at small stimulus sizes, while at large sizes the inhibition was still dominant (in line with ***Figure 3***). In contrast, in non-overlapping receptive field regions, suppression of cortical feedback (***Figure 5***f) resulted in a withdrawal of indirect additional inhibition coming di-synaptically from the cortex, which consequently led to an enhanced response at large stimulus sizes, while at small sizes both the feedforward and feedback inhibitions were less engaged.

Taken together, from the thalamus-centric point of view, our model shows how the corticothalamic loop could significantly contribute to the interactions of cortical visual space representations through their back-projection to the thalamus. In particular, the feedback promotes a unified stimulus representation between cortex and thalamus over short distances: it provides excitation for spatially overlapping receptive fields (***Figure 5***ace), and it imposes a retroaction of the cortical orientation preference bias onto the thalamus (***Figure 4***cde). In contrast, over medium distances, the feedback promotes competition between different stimulus representations: the feedback provides suppression for spatially non-overlapping receptive fields (***Figure 5***bdf), increasing the intrathalamic surround suppression (***Figure 3***).

### The Cortical viewpoint

In the previous sections, we have shown how cortical activity exerts influence over the thalamus through the cortico-thalamic feedback connectivity, and hence recurrently reshapes its own input, and consequently its activity. We will now explore the hypothesis that the thalamo-cortical loop allows the cortex to up- or down-regulate its own functional selectivity in a stimulus-dependent fashion.

#### The loop enhances cortical selectivity to stimulus size and orientation

A series of electrophysiology studies showed that the activity of LGN neurons that further project to V1 is tuned to the size of the stimulus with larger than preferred stimuli suppressing the response of the neurons (***Nolt et al., 2004***; ***Sceniak et al., 2006***), even in the absence of cortical input. The exact mechanisms of such surround suppression in LGN remain unclear, and no computational models offering mechanistic explanations were proposed yet. This surround suppression in LGN is enhanced further in the cortex (cat: ***Hubel and Wiesel, 1965***; ***Sillito et al., 1993***, monkey: ***Sceniak et al., 1999***; ***Angelucci and Bressloff, 2006***). In our model, we found that corticothalamic feedback enhances suppression for non-preferred stimuli both with respect to the size and orientation of the stimulus.

We presented our model full-field drifting gratings varying in orientation and drifting gratings patches varying in size while recording cell responses from a circular region within a cortical ISO-preference domain, in two model configurations: “full”, and “feedforward-only” (***Figure 6***). In this latter case, the cortico-thalamic feedback connections were absent, corresponding to an in-vivo preparation in which the corticothalamic pathway would be inactivated (for example by inactivating V1 Layer 6 corticofugal neurons with muscimol, as in ***Jones et al. (2012***), or using optogenetics, as in ***Hasse and Briggs (2017***)). We found that the presence of cortico-thalamo-cortical feedback was associated with enhanced suppression at stimulus sizes beyond preferred, and at stimulus orientations cross-oriented with the preferred one.

**Figure 6.**
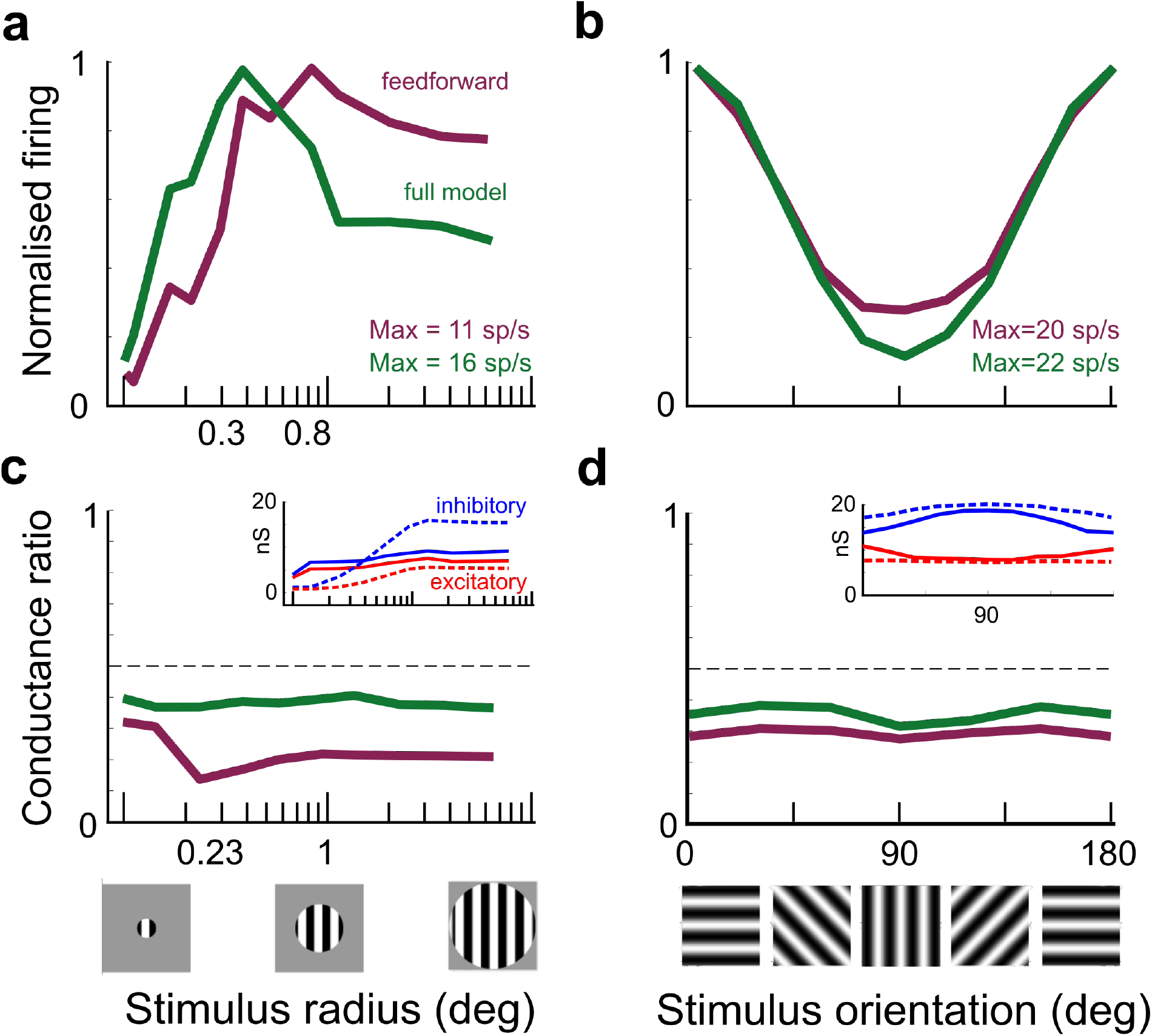
The loop increases size and orientation selectivity by increasing suppression for non-preferred stimuli. (**a**) Min-Max normalized trial-averaged size tuning curves for (n=32) cortical cells. Responses were more suppressed for larger-than-preferred sizes (>1.5 deg) in the full (green) compared to the feedforward-only (purple) model configurations. (**b**) Normalized trial-averaged orientation tuning curves for cortical cells. Responses were lower for stimuli orthogonal to the preferred orientation in the full model (green) vs the feedforward-only (purple) model configurations. (**c**) Excitatory-to-inhibitory size conductance ratio. The closing of loop reduced the relative impact of inhibition, with E/I ratio values closer to 0.5 for the full (green) vs feedforward-only (purple) model configurations. Inset shows the corresponding trial-averaged mean excitatory (red) and inhibitory (blue) synaptic conductance size tuning curves of model V1 cells in the full (solid) vs feedforward-only (dashed) configurations. (**d**) Excitatory-to-inhibitory orientation conductance ratio. Although the presence of the closed loop altered the response for orthogonal stimuli, there was only a minor change in excitatory/inhibitory balance. In the inset, trial-averaged mean excitatory (red) and inhibitory (blue) synaptic conductance orientation tuning curves of model V1 cells.

In order to compare cortical responses in the two configurations, we measured the “feedforward-only” (purple) and “full” (green) trial-averaged normalized firing rates (***Figure 6***ab). For the size tuning protocol (***Figure 6***a), in the “feedforward-only” configuration the suppression index was low (SI: 0.11±0.13), while in the “full” model, the presence of the cortico-thalamo-cortical feedback pathway resulted in a significantly higher values of the suppression index (SI: 0.32±0.12, Welch t-test, p=0.0052). For the orientation tuning protocol (Figure 6b), in the feed-forward-only configuration, the suppression for non-preferred stimuli was significantly lower than in the full model (−8.8%, Welch t-test p=0.0071).

We explored the underlying relationship between excitatory and inhibitory synaptic conductances through size and orientation tuning curves recorded in both model configurations by computing the excitatory-to-inhibitory (E/I) conductance balance. For the size tuning protocol (***Figure 6***c), the cortico-thalamo-cortical loop shifted the conductance regime of excitatory cells from dominant inhibition in the “feedforward-only” configuration (0.23) to a more balanced conductance regime in the “full” model (0.41). For the orientation tuning protocol (***Figure 6***d), the corticothalamic feedback shifted the conductance regime of excitatory cells towards a more balanced (feedforward-only: 0.29; full: 0.37). In the “feedforward-only” configuration, excitatory cells were dominated by inhibitory conductances (insets of ***Figure 6*** c and d, for both types of stimulus variation). Our “full” model results for size tuning of firing rate responses were in agreement with the available cat V1 data (***DeAngelis et al., 1994***, SI: 0.52±0.4). And the radius sizes for the tuned responses of center and surround corresponded to the measures reported in the literature (***Sillito et al., 1993***).

Importantly, the reduction of suppression measured for “feedforward-only” vs “full” configurations was in agreement with the only (to our best knowledge) available cat experimental data about size-tuned cortical responses in the absence of cortical feedback to the thalamus, provided by ***Bolz and Gilbert (1986***). They inactivated Layer 6 of cat V1 with GABA injections, while presenting stimuli varying in size, and reported a reduced suppression index in Layer 4 (SI changed from 0.54 to 0.10). However, Layer 6 is known to also be a source of direct input to Layer 4 (***Douglas and Martin, 1991***), therefore their results could not be unequivocally attributed to the cortico-thalamo-cortical feedback loop, unlike in our model, where the only external source of input to cortical neurons comes from the thalamus.

In summary, our model shows that the cortico-thalamic loop enhances suppression for non-preferred (both in terms of orientation preference and size) stimuli, facilitating (i) competition between local representations of different orientations and, (ii) competition between representations of retinotopically proximate stimuli. These suppressive mechanisms are mediated both by local recurrent cortical circuitry and also long-range lateral connectivity (***Kapadia et al., 2000***; ***Stettler et al., 2002***).

#### The loop enhances competition within the hypercolumn but not in the surround

So far we have studied the cortical response to stimulus size and orientation independently. We thus next proceeded to study whether the cortico-thalamic loop engages the short vs. long-range cortical circuits differentially and whether these lateral interactions depend on the orientation preference of the neurons. To do so, we extended our analysis to also include (i) cells spatially offset from the cortical location retinotopically aligned with the stimulus center, and (ii) cells with cross-oriented preference to that of the stimulus. In the previous section, we found that the corticothalamic feedback facilitates competition between the representation of nearby stimuli through enhanced surround suppression. Here we found that this competition is restricted to the hyper-column aligned with the stimulus center and the strength of these competing influences is independent of the orientation preference of the modulated neurons.

We analyzed two topologically defined domains, “Center” and “Surround” (***Figure 7***a). The first domain contained cortical cells with RF centers aligned with the center of the stimulus and located within a circular area of 700 *μ*m radius, roughly the scale of a hypercolumn. The other domain contained cells with RF centers positioned within the surrounding annulus of the visual space (inner circle of 1 degree radius and outer circle of 1.8 degrees radius), and situated beyond 1000 *μ*m from the cortical position co-aligned with the stimulus center. Both in the center and surround domain, neurons were partitioned into ISO-group containing cells with orientation preference matching the stimulus orientation, and the CROSS-group containing cells with orthogonal preference. By convention, the grating orientation was set at the preferred orientation 0 degrees. In order to compare the responses of all groups, we reported the normalized firing rate in the “full” (green) and “feedforward-only” (purple) model.

**Figure 7.**
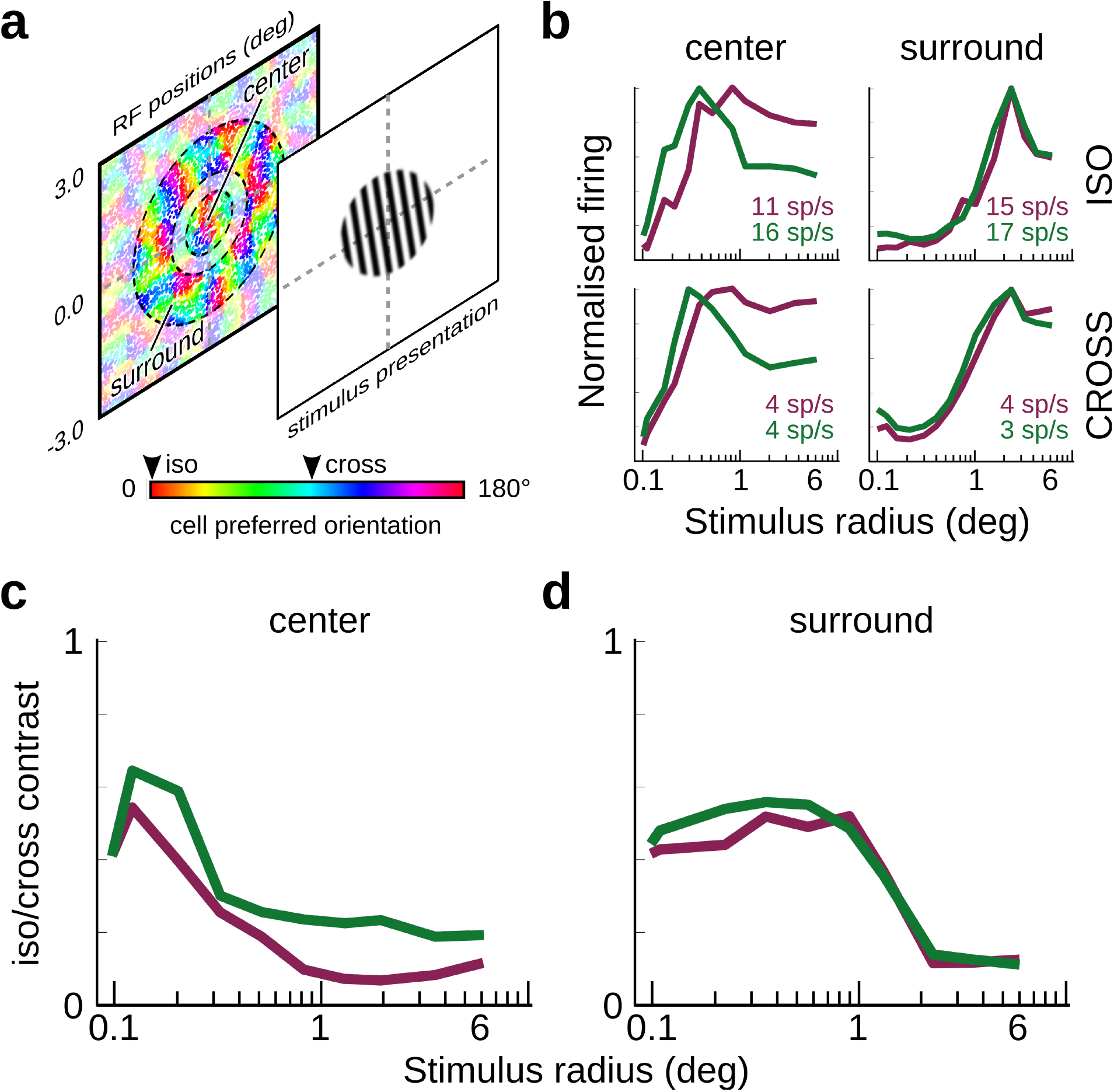
The loop increases the competition between the center and surround, but only within the hypercolumn encoding the stimulus center. (**a**) cells had their RF centers in either a center domain (0.7 degree radius) or a surround annulus domain (from 1.0 to 1.7 degrees). Within each domain, we considered cells having a preferred orientation matching that of the stimulus (0 degrees; ISO group) and cells having the orthogonal preferred orientation (∼90 degrees; CROSS group). (**b**) Normalized tuning curves for all groups in both domains. (Left column) Center-ISO (n=32) and Center-CROSS (n=42) mean responses reached their peak response around 0.3 deg radius and were more suppressed for large sizes (>1.5 deg radius) in both the “full” model (green) compared to the “feedforward-only” configuration (purple). (Right column) No significant reduction was measured for the Surround-ISO (n=561) and Surround-CROSS (n=591) cell groups. Both reached their peak around 2.1 deg radius. (**c**) The orientation map contrast (ratio ISO/CROSS activities) in Center groups is affected differently, depending on the model connectivity configuration. The feedforward-only configuration had significantly lower tuning contrast than the full (t-test, p=0.003). (**d**) In the surround groups, the tuning contrast levels were not significantly different between the feedforward-only and full model configuration.

In the center, the ISO group mean responses were more suppressed for large sizes (>1.5 deg) in the “full” model (green) compared to the “feedforward-only” configuration (purple), as already shown in ***Figure 6***. Interestingly, both neurons in the ISO and CROSS groups exhibited a similar increase in surround suppression in the “full” relative to the “feedforward-only” condition, in line with the fact that the additional surround suppression in the “full” condition is mediated by the LGN neurons that lack (strong) selectivity to the orientation of the stimulus. Furthermore, such change in surround-suppression in the model V1 was absent in the neurons recorded in the surround region.

In order to quantify the competition between ISO and CROSS groups, we looked at the tuning of the orientation map contrast — calculated as the normalized ratio between responses of ISO (*R*_*i*_) and CROSS (*R*_*c*_) orientation preference domains (*R*_*i*_ − *R*_*c*_/*R*_*i*_ + *R*_*c*_). We observe that the corticothalamic loop increased the orientation tuning contrast in neurons located in the center domain (***Figure 7***c), but this change was absent in neurons located in the Surround-domain (***Figure 7***d).

These results showed that the cortico-thalamic loop reinforces locally the competition between ISO-functional domains by sharpening the apparent contrast of the orientation preference map within the hypercolumn retinotopically co-aligned with the RF center. This effect is however absent in neurons recorded in the surrounding region of the cortex, indicating that the cortico-thalamic loop engages only a short-range cortical circuit, in line with the limited retinotopic range of the feedback cortico-thalamic connection.

#### The loop selectively suppresses cortical functional connectivity

In the previous section, we found that the cortico-thalamic loop increased competitive interactions between ISO- and CROSS-oriented groups within a cortical hypercolumn. We then hypothesized that the loop could change cell-to-cell interactions in a stimulus-dependent manner, thus affecting cortical functional connectivity. To test this hypothesis, we characterized cortical functional connectivity as the capacity of one portion of the cortex (i.e. the Center) to alter the cell excitability of another portion of the cortex (i.e. the Surround), with respect to the impact locus of feedforward stimulation. We then inspected how the loop modulates these interactions. In our model, we found that the cortico-thalamic loop induces suppression of the local intracortical cooperative facilitation of ISO-but not CROSS-oriented groups, for larger than preferred stimulus sizes, thus demonstrating the selective influence of the corticothalamic loop on the interactions across visual space.

To characterize functional connectivity with respect to cortical location, cell stimulus feature preferences, and the cortico-thalamic loop, we recorded the spikes, synaptic conductances, and membrane potentials, from ISO- and CROSS-cell groups, in both Center and Surround locations (the same cell group definitions in ***Figure 7***a of the previous section). We then performed a population spike-triggered average (STA) of the synthetic local field potential (sLFP). The sLFP, computed using membrane potential in conjunction with excitatory and inhibitory conductances, produces the currents measured by a virtual electrode (***Destexhe et al., 1998***, see Methods and supplementary figure 3). The sLFP captures not only the balance between conductances but also their effect relative to the state of depolarisation of the local neural population. As such, it is a measure of the ‘effective’ input currents driving the postsynaptic membrane potentials in the surrounding population of neurons (***Destexhe et al., 1998***). By averaging sections of sLFP recorded in one location triggered on spikes emitted by the population of neurons at another location, population-to-population cortical functional interactions can be measured (***Eckhorn et al., 1988***; ***Arieli et al., 1995***; ***Katzner et al., 2009***; ***Nauhaus et al., 2009***; ***Einevoll et al., 2013***; ***Baudot et al., 2013***). The magnitude of the through of such STA of sLFP, is indicative of the strength of depolarizing interactions between the population. We will refer to this quantity as depolarizing interactions index (DII) in the following text. We computed the DII for every combination of cell groups (ISO vs CROSS, ***Figure 8*** c and d vs e and f; and center STA to surround spikes vs surround STA vs center spikes ***Figure 8*** c and e vs d and f), in each cortical location, for each stimulus size (10 patches of drifting gratings of varying size, 12 trials), in the “feedforward-only” and “full” model configurations.

**Figure 8.**
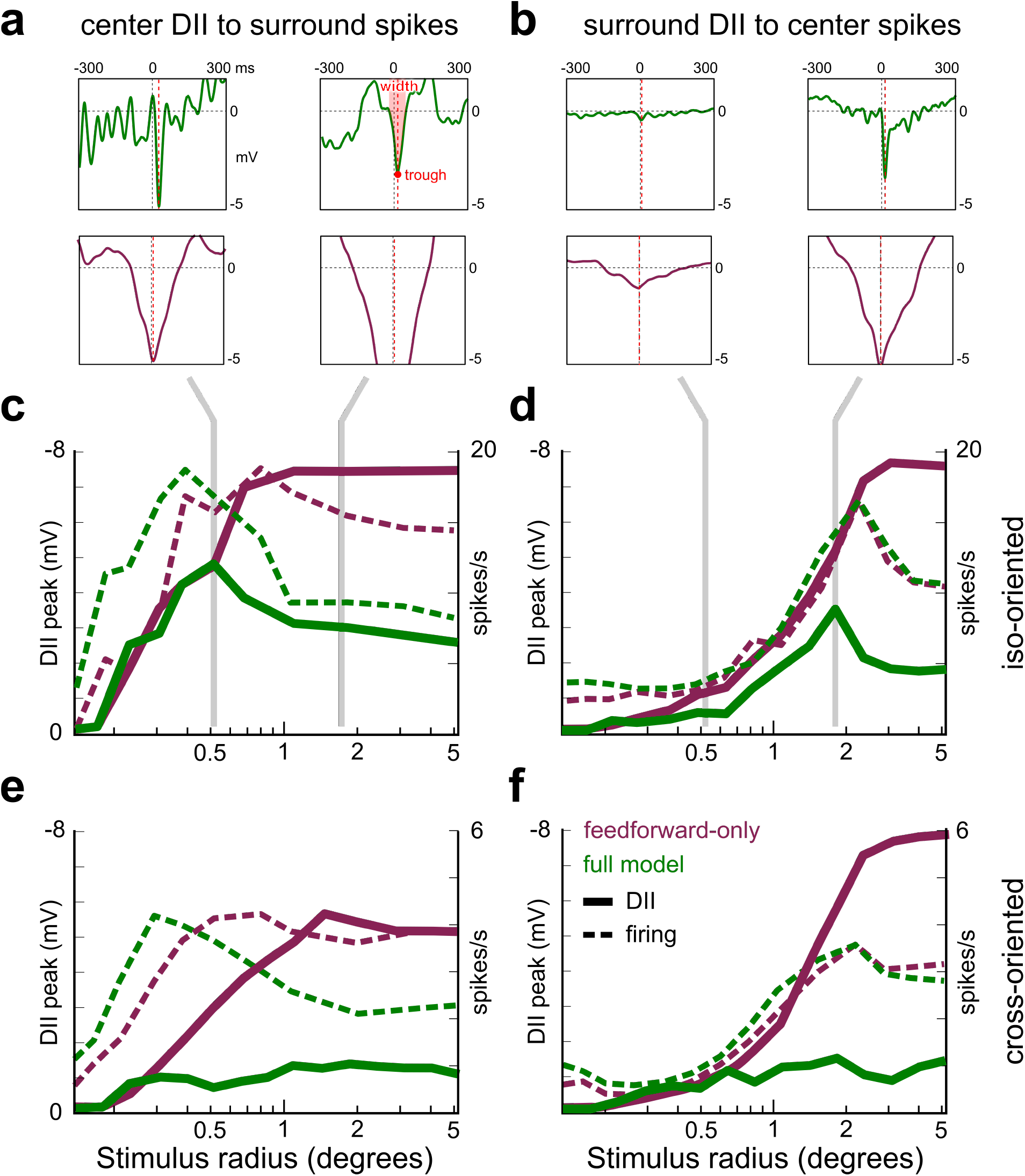
Population spike-triggered average of sLFP amplitude size tuning curves. (**a-b**) Example STA of Center-sLFPs triggered by spikes in the Surround (**a**), and similarly of Surround-sLFPs triggered by spikes in the Center (**b**), measured at preferred (left column) and larger-than-preferred (right column) stimulus sizes, in the feedforward-only (top, green) and full (bottom, purple) configurations. (**c-d**) Size tuning curves of DII (the peak amplitudes of sLFP STAs waveforms) for the center STA to surround spikes and surround STA to center spikes conditions and ISO- and CROSS-oriented cell groups in the cortex (solid, respectively, c: n=38, e: n=16, d: n=31, f: n=21). Corresponding size tuning curves of firing rates are dashed, as in Figure 7. (**c**) Cortical DII peak amplitude (solid curve) of ISO-oriented cells in the Center, triggered by spikes in the Surround. In the “feedforward-only” configuration (purple), the DII is reaching its peak maximum at 0.5 deg of stimulus radius. For larger stimuli, the DII saturates. In the “full” model (green), the DII peak amplitude is overall lower, with a corresponding peak at 0.5 deg stimulus radius, but also remains significantly lower than in the “feedforward” configuration for larger stimuli. The DII tuning roughly followed the firing tuning (dashed). (**d**) The surround DII response of ISO-oriented cells triggered by Center spikes follows a similar trend, with its maximum shifted to larger (2.1 deg radius) stimuli. (**e-f**) The corticothalamic loop only reduces DII amplitudes for CROSS-oriented cells in both the center and surround.

The DII followed quite different trends across configurations and stimulus sizes. To facilitate comprehension and comparison with previous sections, we inverted the sign of sLFP vertical scales in the panels of ***Figure 8***. Overall, there was a clear reduction in the amplitude of the DII in the “full”-loop model configuration relative to the “feed-forward” model configuration (***Figure 8*** cf). In both ISO-group conditions (***Figure 8*** c,d, green lines), the DII tuning curve for the “full”-model configuration exhibited size tuning, and the stimulus size at which the DII and firing rate size tuning curves reached maximum roughly coincided. In contrast, there was an absence of size tuning of the DII size tuning curves in both ISO-group conditions for the “feed-forward” model configuration (***Figure 8*** c,f; purple lines). Interestingly, in both CROSS-group conditions, there was a lack of size tuning in the DII curve for both model configurations (***Figure 8*** e,f).

In summary, the cortico-thalamic loop maintained, relative to the “feed-forward” configuration, the facilitatory interactions between neural populations in the Center and Surround, whose functional preference matched the orientation of the stimulus, but only for stimulus sizes that were confined to CRF (as defined by the peak of the size tuning curve). But, the facilitating interactions were suppressed once stimuli started invading the surround. In contrast, for neurons whose preference did not match the stimulus orientation, a suppression (relative to the “feed-forward” configuration) was found irrespectively of the stimulus size. This demonstrates the potential impact of the cortico-thalamic loop to modulate interactions across cortical and hence visual space in a functionally specific and stimulus-dependent manner. We hypothesize that this behavior could further enhance the role of V1 in the extraction of contours under cluttered conditions.

Taken together, from the cortex-centric point of view in ***Figure 6*** we first showed that the loop increases cortical suppression for larger than preferred stimulus sizes and non-preferred stimulus orientations. Then, in ***Figure 7***, we showed that this extra feedback mediated suppression facilitates competition between neural populations selective to mutually orthogonal orientations, but these competitive effects are confined within the hypercolumn. In this section, we showed that this additional feedback-mediated surround suppression is induced through the reduction of facilitatory lateral interactions between populations of neurons in the center and surround representing similar stimulus orientations.

## Discussion

The early sensory pathways integrate numerous reverberating loops between the cerebral cortex and thalamus (***Jones, 1985***). While the feedforward visual sensory pathway from LGN to V1 has been studied extensively, the understanding of corticothalamic feedback is limited, due to the existence of only sparse structural data (***Sherman and Koch, 1986***; ***Basso et al., 2005***), diversity of experimental conditions (***Ghodrati et al., 2017***), and lack of reproducible functional observations (***Jones, 1985***; ***Usrey and Alitto, 2015***).

These practical hindrances to advance our understanding of the cortico-thalamic loop are further exacerbated by more conceptual issues. A potential risk in the reductionist practice of disabling (e.g. pharmacologically, through lesioning, or otherwise) an inherently recurrent system at a specific point in the loop, is to attribute any resulting loss of function to the sole locus of the intervention without taking into account the distributed changes occurring concurrently elsewhere. Such reasoning is highly problematic since the function under study can emerge through interactions between many parts of the global neural system (here the early visual system), and local elimination of any of these contributing components could lead to the global function disappearance. Computational models provide new probes and perturbation paradigms that can be applied in any possible location in the detailed virtual architecture of the system under study.

Data-driven simulations concur to a better understanding of the emergent properties through the explicit construction of the recurrence built in the model itself. Indeed, in this study, we have applied such a computational approach to the functional dissection of the role of feedback between V1 and LGN. Our simulations show for the first time that the cortico-thalamic loop yields phenomena that do not arise in LGN or V1 alone, and that the competitive interactions that are typically attributed to cortico-cortical communication can also arise or be further amplified through the cortico-thalamic loop.

The simulations recapitulate - within a single model - experimental observations in cat visual system across several stimulus features and circuit manipulation protocols. The model suggests mechanistic scenarios by which the corticothalamic loop enhances stimulus features already represented by the thalamic circuits (***Figure 3***a), while also strengthening locally functional contrast in the cortically encoded feature maps (***Figure 4***cd, ***Figure 5***cd). The model also describes the mechanism by which the cortico-thalamo-cortical loop supports increased stimulus feature selectivity by increasing the relative difference between levels of activation of local neural populations representing orthogonal stimulus orientations (***Figure 6*** and ***Figure 8***). Particularly, the model indicates that the cortico-thalamic loop may enhance competition between feature-selective domains coexisting within the same hypercolumn but not beyond (***Figure 7***). The model suggests also that the loop selectively suppresses cooperative facilitation between ISO-preference cortical domains (***Figure 8***).

Several previous modeling works already explored corticothalamic feedback. Most of these studies focused on the involvement of the thalamocortical loop in the size-tuning of LGN and V1 neurons. First ***Bonin et al. (2005***) highlighted the limitations of a simple Difference-of-Gaussians (DoG) model to reproduce LGN responses in intact (closed-loop) preparations. They showed how extending the DoG model with a suppressive term, made of a feedforward cascade of linear filters, could offer a better approximation of the early visual system. This model, while only reproducing contrast and size tuning results, still has the important benefit of being linear, without requiring any recurrence. Later, ***Einevoll and Plesser (2012***) extended the Difference of Gaussian model by incorporating terms representing the corticothalamic feedback from a set of idealized orientation-selective cortical cells. Their detailed model reproduced the responses of LGN cells to several stimuli, including size-varying patches of drifting gratings, replacing the suppressive linear filters with an idealized corticothalamic kernel. Recently, ***Martínez-Cañada et al. (2018***) took a more mechanistic approach, simulating a thalamocortical spiking network with LGN relay cells and interneurons, and cortical cells. The network input was modeled as two DoG filter arrays of antagonistic center-surround (ON- and OFF-center) arrangement, as in our model. The thalamic afferents to the cortex were arranged in such a way as to induce phase-opponent Gabor-like ON- and OFF-simple cells, also in line with our model. But, in contrast to our present study, the cortical response was fed back to thalamic relay cells along connectivity configurations dependent on the cortical phase. Two feedback connectivity schemes were tested. One phase-matched scheme, in which ON-center simple cortical cells projected to ON-center thalamic cells (and OFF-center simple cortical cells to OFF-center thalamic cells), and a phase-reversed scheme, in which ON-center simple cortical cells projected to OFF-center thalamic cells (and vice-versa). In accordance with the available literature, they found that a phase-reversed feedback kernel provided an increased center-surround antagonism in LGN responses to patch gratings. In the present study, we show that the assumption of such phase-specific feedback connectivity is not necessary for explaining the center-surround antagonism in LGN responses. This kernel-based approach was also recently used to explain the impact of cortical feedback on LGN representation in a mouse preparation (***Born et al., 2021***). Interestingly, in this work, a wide inhibitory feedback coupling kernel was used to reproduce feedback-enhanced surround suppression and sharpening of LGN receptive fields. Our model incorporates previously characterized (***Deleuze and Huguenard, 2006***) spread of PGN activity, giving biological credence to the cortical-driven source of widespread inhibition to LGN, without the need for phase-specific cortico-thalamic connections.

While our model offers a mechanistic explanation for several important computations taking place in LGN and cortex, the broader purpose of these computations remains to be elucidated. Here, we consider that the selectivity required to categorize visual stimuli and the flexibility to cope with the combinatorial explosion of possible configurations of visual features need to be faced in the early stages of visual processing (***Barlow and Levick, 1969***). Both goals can be achieved in a system that supports selectivity emergence through recurrent competition (***Edelman, 1993***), and flexibility through synergistic interactions (***Kauffman et al., 1995***). We propose that the corticothalamic loop enhances both competition and synergy, thanks to a known connectivity motif, at the heart of our model: focused direct excitation, surrounded by wide indirect inhibition. The crux here is that the interaction domain in the model is not space itself but the cortical feature preference map. Competition is limited to the hypercolumn scale, i.e. covering in cortical space the representation of one point in visual space through all possible functional filters (defined here by the local topology of the orientation map). The synergy between stimulus representations is mediated by corticothalamic direct narrow excitation, where each cortical representation projects to the thalamus its own attuned selectivity, to reverberate again to the cortex (***Alonso et al., 1996***; ***Andolina et al., 2007***; ***Béhuret et al., 2013***; ***Bijanzadeh et al., 2018***). This reverberation competition principle is ubiquitous and has been proposed in other sensory systems (auditory system: ***Suga et al., 1997***; ***Suga and Ma, 2003***; somatosensory system: ***Ghazanfar et al., 2001***; ***Li and Ebner, 2007***; ***Temereanca and Simons, 2004***). Similarly, the just as ubiquitous competition between stimulus representations is mediated by corticothalamic indirect broader inhibition, where each cortical representation suppresses its topological surround, through the thalamic reticular nucleus, source of thalamic inhibition (for the visual system: ***Tsumoto et al., 1978***; ***Funke and Eysel, 1998***; ***Sillito and Jones, 2002***; see for the auditory system: ***Suga et al., 2000***; ***Yan and Suga, 1996***; for the somatosensory system: ***Lam and Sherman, 2005***). The structural-functional model of the early visual system presented here integrates such an unprecedented breath and detail of these key anatomical and functional features. Hence, it constitutes a novel versatile computational toolbox for testing alternative hypotheses on the function of the cortico-thalamic loop, that cannot be addressed through direct physiological or pharmacological lesion experiments.

## Methods and Materials

Here we provide the description of the principles used to construct the simulated cortical and thalamic regions, the various connection pathways within and between these regions, and the cellular and synaptic properties. We also detail the experimental protocols, the procedures for the collection and analysis of data, and the software stack used to develop and simulate the model. The full code listing and parameters are available on the project page on GitHub (https://github.com/dguarino/T2).

### Model structure

Our model was based on a previous model of V1 (***Taylor et al., 2021***; ***Antolík et al., 2019***) that has been modified to focus on the thalamus and the corticothalamic feedback loop. It consists of: (i) a region of visual field, (ii) two retinotopically congruent regions of the dorsal thalamus, namely the lateral geniculate nucleus (LGN) and the peri-geniculate nucleus (PGN), and (iii) a retinotopically congruent region of the cat primary visual cortex (V1). The model covers 7×7 degrees of visual field within 5 degrees of eccentricity around the area centralis. This corresponds in the cat to 1.4 mm2 of LGN and PGN tissue (***Sanderson, 1971***), and to a 3.5×3.5 mm patch of primary visual cortex.

### RETINA

We use a simple model of the transduction from luminance stimuli to retinal ganglion cell (RGC) spikes that includes a phenomenological model of RGC receptive field (as in ***Rodieck, 1965***; ***Cai et al., 1997***). This model has two space-time separable components, one for the center and one for the surround of the RGC receptive field (RF) response:

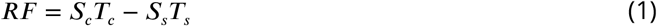

The spatial components was modeled as a Gaussian functions of the visual stimulus position (x), according to the model of ***Rodieck (1965***):

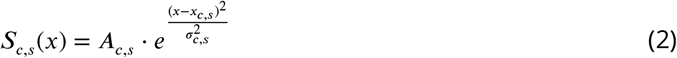

with *A* being the amplitude of response, and *σ* the radius of the RF component. These parameters were chosen according to the literature for cat RGCs at ∼ 5 degrees of eccentricity from the area centralis, to be *σ*_*c*_ = 0.2*deg, σ*_*s*_ = 0.7*deg* (***Linsenmeier et al., 1982***; ***Marrocco et al., 1982***) and *A*_*c*_ = 1.0, *A*_*s*_ = 0.05 (***Cai et al., 1997***). The time components are modeled as sums of Gamma functions (***Cai et al., 1997***):

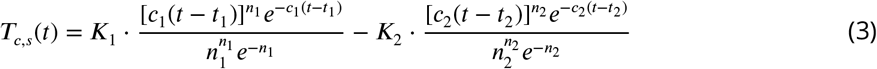

where the parameters were taken from ***Nirenberg et al. (2010***) (*K*_1_ = 1.05, *K*_2_ = 0.7, *c*_1_ = 0.14, *c*_2_ = 0.12, *t*_1_ = −6.0, *t*_2_ = −6.0, *n*_1_ = 7.0, *n*_2_ = 8.0).

The visual transduction, for each RGC, is computed as follows. A *view* of the stimulus, a set of luminance values pertaining to a cell RF area, is convoluted with a RF:

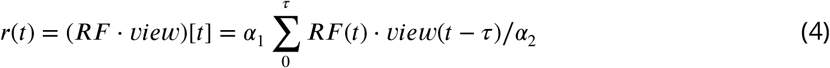

where both RF and view are 3D arrays (2D for the space component and a third dimension for the time component), *τ* = 7*ms* is the duration of the RF temporal component, and *α*_1_ = 380000 and *α*_2_ = 150, are dimensionless linear luminance gain parameters. This response *r*(*t*) represents the absolute mean luminance convolution of RF kernel and stimulus view, therefore it is unbound. We adopted an additional saturation term, similar to Bonin et al. 2005, to saturate the luminance response with a contrast response, obtained by convolving the standard deviation of the spatial component (here simply represented by the variable *x*) of the stimulus *view*(*t*) with the response

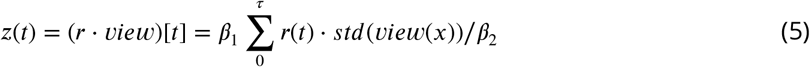

where *β*_1_ = 200000 and *β*_2_ = 0.00001, are dimensionless linear contrast gain parameters. This saturation term represents an abstraction of the RGC firing rate, as if already weighted by the triadic dendro-dendritic contribution coming from LGN interneurons (as suggested by ***Sherman, 2004***), which we did not model explicitly.

The sum of response *r*(*t*) and saturation term *z*(*t*), in units of luminance (cd/m^2^) assuming the kernel values as dimensionless, is injected as a current (nA) in an Integrate-and-Fire model, together with an amount of white noise used to mimic the response variability and average firing rate of RGCs (for On- and Off-center cells, see table 1). The parameters for luminance, contrast saturation term, and noise were chosen to match the literature available for luminance (***Barlow and Levick, 1969***) and contrast (***Derrington and Lennie, 1982***) of RGC responses (***Rathbun et al., 2010***) and S-potential inputs to LGN (***Weyand, 2007***), see supplementary figures 1 and 2.

### LATERAL GENICULATE AND PERIGENICULATE NUCLEI

The lateral geniculate nucleus (LGN) of cats has three laminae of alternating ocularity (***Sherman and Guillery, 2006***, p.48). We modeled one superficial lamina A, containing X cells, further divided into X-On and X-Off cell groups according to their contrast phase preference (***Enroth-Cugell et al., 1974***). We chose to not model the Y pathway of cat since its cells present a non-linear spatio-temporal RF, and we chose to restrict ourselves to the X pathway having a linear response to stimuli (***Sherman and Guillery, 2006***, p.48). The total number of neurons in the superficial LGN lamina is approximately 450000 (***Sherman and Koch, 1986***; ***Budd, 2004***) which, for a total area of 16 mm^2^ and a layer width of 500 *μ*m (***Sanderson, 1971***) gives approximately 56000 cells/mm^3^. Using a ratio of principal cells to interneurons in the range of 3:1 to 6:1 (***Tömböl, 1967***), and considering an estimate of approximately 45000 principal cells per mm^3^ (***Sanderson, 1971***), the number of LGN principal cells would lie between 7500 and 15000 cells per mm^3^. In our model, we adopted the simplification of dropping the third dimension and each layer is two-dimensional only. Given a magnification factor of 0.2 mm2 per visual deg^2^, the amount of principal cells would be ∼9000 cells per mm^2^, or ∼1800 per deg^2^. For practical reasons of simulation time, we down-sampled it to 100 principal cells per deg^2^, with a total 12800 principal cells in the LGN, divided into two 6400 sets for On- and Off-center cells. We applied the same reasoning for PGN, using the same cell density and magnification factor (***Sanderson, 1971***), given the receptive field size overlap observed in the literature (***Lam and Sherman, 2005***; ***Soto-Sánchez et al., 2017***).

### PRIMARY VISUAL CORTEX

The cortex corresponds to a 3.5×3.5 mm patch of cat primary visual cortex. It contains 10800 excitatory and 2700 inhibitory neurons, in the ratio 4:1 (***Beaulieu and Colonnier, 1989***; ***Markram et al., 2004***), and 10 million synapses, with a significant downsampling (∼10%) of the actual density of neurons present in a corresponding portion of cat cortex (***Beaulieu and Colonnier, 1989***) to make the simulations computationally feasible. For further details on the cortical structure, please refer to ***Antolík et al. (2019***).

### Model neurons

All excitatory and inhibitory neurons are modeled as point-like spiking neurons. All sub-cortical neurons are conductance-based Leaky Integrate-and-Fire (LIF, ***Lapique, 1907***; ***Dayan and Abbott, 2005***). We adapted LIF unit parameters for the different cell types from in-vivo and in-vitro measurements of neurons in the cat visual system and classically modeled in the literature (***Freed and Sterling, 1988***; ***Worgotter and Koch, 1991***; ***Destexhe et al., 1996, 1998***; ***Huertas et al., 2005***; ***Budd et al., 2010***), and the neuroelectro db (neuroelectro.org). The neurons in cortex are modeled as Adaptive Exponential Integrate-and-Fire units (***Brette and Gerstner, 2005***) and their parameters are detailed in table 1 (see ***Antolík et al., 2019*** for motivation and justification). All neuron model details may also be found in the online repository for the project.

### Connectivity

In constructing the model, we have included realistic network properties, such as the spread and relative proportions of the various sets of connections composing the intra-, inter-thalamic, and thalamocortical circuitry. The synaptic connections are probabilistically drawn with replacement, and number of synapses given in the table above. However, we allowed formation of multiple synapses between neurons, hence the exact number of connections between neurons is variable. The extents of axonal and dendritic arborization reported in the literature usually refer to the terminal extents of labeled cells; we took these measures as representing three standard deviations (3*σ*) of the Gaussian distribution of connection probability. Another important consideration in designing the connectivity is the reliability of synaptic transmission. Both in cortex and thalamus, it has been shown that a small fraction of presynaptic action potentials succeeds in evoking postsynaptic potentials (***Allen and Stevens, 1994***; ***Stratford et al., 1996***; ***Weyand, 2007***). However, it would be computationally expensive to model explicitly synaptic failures, therefore we have decided to account for synaptic transmission failures in the number of simulated synapses per neuron, adopting a 10% factor (unless otherwise stated in the following text) with respect to the input counts reported in the literature.

### THALAMIC CONNECTIONS

For all intra-thalamic connections, LGN-to-PGN and PGN-to-LGN, we used a zero-mean Gaussian probability distribution:

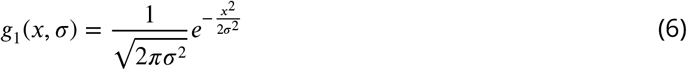

with standard deviation as mentioned in the paragraph above. The total number of synapses per LGN cell is reported to be between 5000 and 8000 (***Sherman and Guillery, 2006***; ***Murphy et al., 2000***; ***Sillito and Jones, 2002***). Of the total, 16% of these connections are from RGCs. ***Sherman and Guillery (2006***) reported that the majority these input cells are not able to trigger an action potential in the postsynaptc LGN target and suggests that 2-5 individual input coincidence is required. In spite of this wiring constraint, we further simplified our model by simulating an simplified monosynaptic connection between an RGC and one LGN principal cell. Then, 36% come from PGN (therefore 1800-2880, mean ∼2200, downsampled to 220), 5% from interneurons (we do not model them, see above). The total number of synapses per PGN cell is reported to be around 6000 (***Golshani et al., 2001***; ***Sillito and Jones, 2002*** p.1659: 6771±1018) of which: 20% is from LGN (therefore ∼1200, downsampled to 120), 15% form other PGN cells (therefore ∼900, downsampled to 90). Measures of the mean arborization distance of LGN axons into PGN are obtained from both morphological (***Friedlander et al., 1979***) and functional (***Lam and Sherman, 2005***) studies, and report radius range between 50 and 150 *μ*m. For our Gaussian distributions, we used a 3*σ* radius of 75 *μ*m. Morphological studies (***Cucchiaro et al., 1991***; ***Sanchez-Vives and McCormick, 1997***; ***FitzGibbon, 2000***; ***Fitzgibbon, 2006***) give a 100-300 *μ*m radius for PGN to PGN connections, either dendrodendritic or axon collaterals. We used 70 μm (3*σ*). Morphological (***Cucchiaro et al., 1991***; ***Cox et al., 1996***; ***Sanchez-Vives and McCormick, 1997***) and functional (***Lam and Sherman, 2005***) studies give 150-300 *μ*m radius for the axonal arborizations of PGN to LGN. We adopted 150 *μ*m radius (3*σ*).

### THALAMO-CORTICAL CONNECTIONS

According to the literature (***Sherman and Guillery, 2006***; ***Budd, 2004***), each cortical neuron receives connections from at least 450 LGN cells (we downsampled it to 45). Each neuron in V1 received connections from both On- and Off-center LGN populations. The spatial thalamo-cortical connectivity was determined by a Gabor distribution, as reported by ***Troyer et al. (1998***):

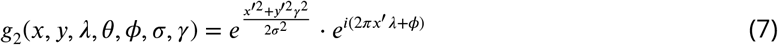

where *x*′ = *xcosθ* + *ysinθ* and *y*′ = −*xsinθ* + *ycosθ*. For individual neurons the orientation *θ*, phase *ϕ*, size *σ*, frequency *λ*, and aspect ratio *γ* of the Gabor distribution were selected as follows. To obtain a functional organization in the cortex, we pre-computed an orientation map corresponding to 3.5×3.5 mm of cortical area (***Antolík and Bednar, 2011***), and this map was used to assign an orientation preference *θ* to each cortical neuron. The phase *ϕ* of the Gabor distribution was randomly assigned ***Ziskind (2013***). We set to constant values the remaining parameters, following the average values reported by Jones and Palmer 1987 for cat V1 RFs located in the parafoveal area (size *σ*=0.25 degrees of visual field, the spatial frequency *λ*=0.8 Hz and the aspect ratio =0.57).

### INTRACORTICAL CONNECTIONS

According to ***Beaulieu and Colonnier (1985***), the number of synaptic inputs per single neuron in cat V1 is 5800. A large portion of these connections comes from other cortical areas ***Budd and Kisvárday (2012***), with recent estimates for the cat, reporting a 76% of them coming from outside V1 (***Stepanyants et al., 2008***). In our model, cortical synapses were formed taking into account the proportion of excitatory and inhibitory cortical cell type densities, the average number of synaptic inputs, the proportion of extra-area input and the failure rates of synaptic transmission. In total, each excitatory cortical cell in our model receive 800 synaptic inputs, while inhibitory neurons receive 520 inputs, to account for their smaller size. The probability of connection between two neurons in the network was distance-dependent, based on Gaussian decay. Cortico-cortical connectivity was determined by the zero-mean hyperbolic probability density function (pdf):

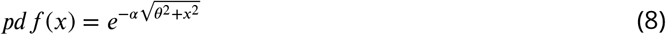

with *α* being a distance parameter, and the other parameters as above. The parameters for this pdf incorporated four known principles: (i) connection probability decays with increasing cortical distance between neurons (Budd and Kisvarday 2001, Stepanyants et al. 2008); (ii) connections have a functionally specific bias (Buzas et al. 2006, Ko et al. 2011); (iii) excitatory neurons have the weak tendency towards connecting nearby neurons of similar receptive field properties, and this tendency increases for more distant post-synaptic neurons (Buzas et al. 2006); (iv) the anti-phase relationship between excitatory and inhibitory conductances in cat V1 simple cells suggests a push-pull connectivity, at least in layer 4 (Hirsch et al. 2003, Lauritzen and Miller 2003, Monier et al, 2006; Baudot et al. 2013).

### CORTICOTHALAMIC CONNECTIONS

Of the total 5000-8000 synapses per LGN cell (Sherman and Guillery 2001, Murphy et al. 2000, Sillito and Jones 2002) 44% come from cortex (therefore 2200-3520, with a mean of 2800, that we down-sampled to 280). The number of synapses per PGN cell coming from the cortex is 60% (therefore ∼3600) but, in this case, a different downsampling reasoning was applied. The feedback connections from primary visual cortex to PGN have been measured only by few morphological studies and reported to form en-passant boutons, in the order of tens, within a radius below 25 *μ*m (Ahlsen and Lindstrom 1983, Boyapati and Henry 1984, FitzGibbon et al. 1999, FitzGibbon 2000). However, PGN cells form a dense dendro-dendritic network that is capable of spreading excitation in a radius of ∼100 *μ*m (Deleuze and Huguenard 2006). For the sake of simplicity, we downsampled the number of cortical synapses onto PGN cells to 40, and we used a radius of 90 μm (3*σ*), that we chose through a parameter search strategy (see below). Cortical inputs to LGN have been measured mainly by morphological studies (Murphy et al. 1999, FitzGibbon et al. 1999, daCosta and Martin 2009) and reported to have irregular circular shapes with apical radius varying widely between 50 and 100 *μ*m. We used a 60 *μ*m radius (3*σ*). We drew all cortico-thalamic connections probabilistically from a zero-mean Gaussian distribution (as in eq. 1, above), in spite of a recently proposed Gabor-shaped distribution of cortico-thalamic connection (Jones et al. 2012): the reason was that the evidence for a Gabor distribution is still very sparse and linked to the presence of extra-striate signals that we did not model.

### Synapses

In the conductance-based neuron models we adopted, all synaptic connections are collapsed into a single synapse mechanism under an assumption of linear summation. For the change in conductance caused by presynaptic events, we adopted a step increase followed by an exponential decay of the postsynaptic conductance change. Retino-geniculate synaptic strength has been measured in-vitro (Blitz and Regehr 2003, 2005, Lam and Sherman 2005) giving an overall change in conductance of 10-40 nS mediated by AMPA synapses; we used 6 nS with 1.5 ms decay time constant for the excitatory synaptic conductance (see table 1). This value is below the experimentally measured ones, because higher values led to excessive network activity. LGN to PGN synaptic strength, mediated by AMPA synapses, has been measured by Liu et al. 2001 on a per synapse basis leading to a value of 1.914±1.814 nS, we used 1.5 nS, with 1.5 ms decay time constant. Recurrent PGN synaptic strength, mediated by GABA synapses, was reported to be in the range 0.3-5 nS (Ulrich and Huguenard 1996); we used a lower 0.1 nS with 5 ms decay time constant for the inhibitory synaptic conductance. The synaptic conductance of inhibitory connections from PGN to LGN, mediated by GABA synapses, has been reported to be in the range 0.4-0.6 nS (Sanchez-Vivez et al. 1997, Lam and Sherman 2005), we used 0.5 nS with 5 ms decay time constant. In cortex postsynaptic conductance change is modelled in the same way. The strength of LGN to V1 synapses, mediated by AMPA, has been measured by Cruikshank et al. 2007 in the 0.7-2.0 nS range. Accordingly, we used 2 nS with a 7.8 ms decay time constant. The cortical feedback synaptic efficacy onto LGN cells is mediated by AMPA synapses, with varying measures in the mouse: 0.128±0.047 nS in (Golshani et al. 2001), 2-8 nS in (Lam and Sherman 2013). We opted for 0.2 nS with 1.5 ms decay time constant. The cortical feedback onto PGN cells has been measured also in the mouse to be 0.400±0.257 nS (Golshani et al. 2001, Liu et al. 2001); we used 0.4 nS with 1.5 ms decay time constant of the excitatory synaptic conductance.

### Delays

The transmission delays between processing layers we adopted were taken from several converging experimental reports: 1 ms from LGN to PGN, 1 ms PGN to PGN (Rogala et al. 2013), and 1 ms PGN to LGN (Lindstrom 1982, Funke and Eysel 1998, Rogala et al. 2013). The transmission delay between LGN layers to the cortex is 2 ms (Lindstrom 1982). While feedback transmission delays between cortex and LGN or PGN are between 3 and 5 ms (Lindstrom 1982, Budd 2008, Rogala et al. 2013).

### Inactivation protocols

Several virtual inactivation experiments were conducted, to compare the model’s output with experimental results from the literature. In order to replicate the in-vivo experimental conditions as closely as possible in-silico, we used three main procedures: (i) reproducing the intact brain (“full” configuration), (ii) inactivations of small portion of cortex (“variable feedback” configuration), (iii) inactivation of only the feedback connections (“feedforward-only” configuration), see figure 1 for a summary. In configuration (ii), a point in cortex was chosen having the same (or normal) orientation with respect to the stimulus. To mimic activity suppression (for instance by muscimol injection), a negative intracellular current, strong enough to prevent reaching spike threshold, was injected into all neurons located within a circular area of variable size (300 and 600 *μ*m, corresponding to 0.3 and 0.6 degrees of visual field), such as to simulate various scales of postsynaptic blocker diffusion. Jones et al. 2012 reported an area of cortical focal iontophoretic inactivation of approximately 600 *μ*m, corresponding to 0.5 degrees considering V1 magnification factor around the area centralis. The relative spatial position of cortical and thalamic receptive fields were also taken into account, identifying overlapping and non-overlapping receptive fields. For the inactivation of cortical non-overlapping areas, the authors reported a thalamic distance between receptive field centers of approximately 2.5 degrees of visual field, that we matched in our model. We selected a circular patch of cortical excitatory units and directly injected a current of −0.5 nA in each simulated cell, resulting in a hyperpolarization that prevented cortical cell response.

### Stimuli

We performed our simulations using clips of visual stimuli (graded in contrast and luminance) specific of each associated experimental protocol. Each stimulus sequence consisted of a series of visual stimuli of variable contrast which were interleaved with the 150 ms presentation of a full field of uniform luminance (50 cd/m2). Each visual stimulus was described, at any given point in space and time, by a single number representing its contrast (relative to the mean luminance of the uniform field), ranging from −1 (dark) to 1 (maximal brightness), with 0 being the chosen background luminance. Six types of stimulus have been used in this work: (a) changes of ambient mean luminance, (b) changes of contrast level in a sinusoidal drifting grating (DG), (c) changes of spatial frequency of DG, (d) changes of temporal frequency of a DG, (e) changes of size of a DG patch, (f) changes of orientation of a DG. The model responses have been tuned using iteratively all stimuli. Stimuli (c), (d), (e), (f) were used to test the model in both control (i) and altered (ii) configurations, while stimulus (e) was also tested with partially inactivated cortex (iii). All stimuli were created using the Mozaik framework (Antolik and Davison 2013, see below for details).

### LUMINANCE

The model was stimulated with five levels of ambient luminance: complete darkness, 0.085, 0.85, 8.5, 85 cd/m2, as in Barlow Levick 1969, and Papaioannou and White 1972. The average (background) level of luminance for all other simulated experiments was 30 cd/m2.

### CONTRAST

We used Michelson contrast, as the difference between the cd/m2 luminance peak (Lmax) and trough (Lmin) values, divided by twice the average (background) luminance, C = (Lmax - Lmin) / Lmax + Lmin). Being normalised by the background luminance, the contrast is usually expressed as percentage. Nine contrast levels were used: 0, 2, 4, 8, 18, 36, 50, 80, 100%, as in Kaplan et al. 1987.The contrast level of all the following experiments (c-f) was fixed at 80%.

### SPATIAL FREQUENCY

Patterns of full-field sinusoidal drifting gratings were supplied to the model. Ten different frequencies were chosen for the sinusoidal gratings: 0.07, 0.1, 0.2, 0.3, 0.5, 0.8, 1, 1.5, 2, 8 cycles/degree (with temporal frequency of 2 Hz), as in Bisti et al. 1977, Maunsell et al. 1999.

### TEMPORAL FREQUENCY

Patterns of full-field sinusoidal drifting gratings were supplied to the model. Ten different temporal frequencies: 0.05, 0.2, 1.2, 3.0, 6.4, 8, 12, 30 Hz were chosen for the drifting movement of the gratings (having 0.5 cycles/degree of spatial frequency), as in Marrocco and McClurkin 1985, Marrocco et al. 1996, Alitto and Usrey 2004.

### ORIENTATION

Patterns of full-field sinusoidal drifting gratings were supplied to the model, with eight different orientations uniformly chosen between 0 and 90 degrees (having 0.5 cycles/degree of spatial frequency and 2.0 Hz of temporal frequency), as in Daniels et al. 1977, Vidyasagar and Urbas 1982.

### SIZE

Patterns differing in the radius of a circular patch of sinusoidal (with 0.5 cycles/degree frequency) drifting (8 Hz) gratings were supplied to the model. Ten radiuses were chosen: 0.125, 0.19, 0.29, 0.44, 0.67, 1.02, 1.55, 2.36, 3.59, 5.46 degrees, as in Murphy et al. 1987, Sillito and Jones 2002, Deangelis et al. 1994, Jones et al. 2012.

### Iterative procedure to find a single parameter set

We systematically presented the stimuli described above to our model and heuristically found a single parameter set that fitted qualitatively and quantitatively (see next section for the analysis). To assess model stability, we then searched the parameter space around two crucial parameters: the arborisation extent (*σ*) of LGN→PGN and PGN→LGN for the feedforward-only configuration, together with V1→PGN for the “full” and “variable feedback” configurations. As a guide through the parameter search, we took as reference parameter the index of end-inhibition, for the LGN-PGN interactions (supplementary figure 4a), and the slope of the regression performed on the mean percentage change as in figure 4 (supplementary figure 4b).

### Simulation recording and Analysis

We presented the above stimuli to the model for several trials (varying from 6 to 12, depending on the simulated clip duration, which in turn depended on the type of stimulus and reference experimental conditions). We recorded the spike timing and synaptic conductances of units from the LGN, PGN, and V1 populations, chosen either randomly or selectively depending on the type of experiment. For the experiments presenting stimuli (a-f) with configurations (i-ii), we recorded randomly from ∼100 cells in both LGN and PGN. For the experiments presenting stimulus (e) with configurations (i-iii) we also recorded from a grid of 20×20 cells (100 *μ*m side, spacing 5 *μ*m) in both LGN and PGN, centred on the (0,0) coordinate of visual space.

### TRIAL AVERAGED FIRING RATE

The first 100 ms from stimulus onset were discarded in order to get rid of the onset flash effect. Then the firing rate was calculated as the average number of spikes over trials per neuron for each stimulus. When a population measure is provided in the text, the average over neurons was also performed.

### LOW-PASS INDEX

Used by Kimura et al. 2013 to describe the degree of spatial frequency tuning of firing rates to low spatial frequencies. It is computed as the ratio of the response magnitude at the lowest spatial frequency eliciting a response to that at maximal response.

### END-INHIBITION INDEX

Used by Murphy and Sillito 1987 to describe the degree of size tuning in LGN cells. It is given by the percentage difference between the peak and the plateau of the size tuning curve, divided into bins from 1 (low suppression) to 10 (high).

### SUPPRESSION INDEX

A measure similar to the above end-inhibition index, computed as follows: SI=1-Response to large gratingsResponse to preferred size grating

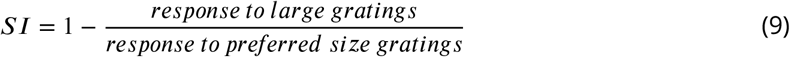

### ORIENTATION BIAS

Used by Vidyasagar and Urbas 1982 to describe the degree of orientation tuning of LGN cells. It is defined as the ratio of the peak response to the preferred orientation to that in the non-preferred orientation for each cell. The results for all cells are grouped into bins from 1 (low bias) to 10 (high).

### SIZE TUNING COMPARISON

As in Jones et al. 2012, after the trial averaged firing rate was computed, only LGN cells showing a significant (p<0.05) change in response were chosen for further analysis. To compare the responses of overlapping and non-overlapping, control and cortex local inactivation, we identified three groups of cells by automatically selecting for each unit the smallest radius eliciting the smallest response, the peak response, and an average of the responses to large radiuses. The comparisons of these responses were done using a non-parametric two-tailed test (Wilcoxon, see below).

### LOCAL FIELD POTENTIAL

Excitatory and inhibitory conductances, and membrane potentials were recorded for all cells within a radius of 3mm from the center of the network (aligned to the presented stimulus center). The transmembrane current of each cell was computed according to the Ohm law:

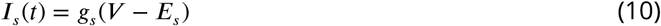

where *g*_*s*_ and *E*_*s*_ denote the conductance and equilibrium potential for each of the two synaptic types (see above Table 1). A biophysical forward-modelling scheme for the LFP was adapted from Einevoll et al. 2013. This method is known to approximate the extracellular electrical potentials generated by cellular transmembrane currents. Its formula is:

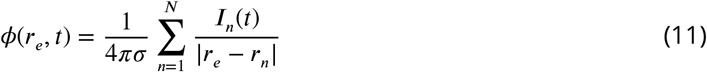

where *I*_*n*_(*t*) denotes the transmembrane current of cell *n* at position *r*_*n*_, *r*_*e*_ is the position of the electrode tip, the sum includes *N* recorded cells, and *σ* is the extracellular conductivity (parameter fixed at 0.1 S/m, according to Dobiszewski et al. 2012). Note that although this method has been developed to synthetize an LFP signal from a multicompartmental model, here we are interested not only at the balance of excitation and inhibition with respect to the membrane potential but also at the effect of distance between contributions. Therefore our use of the formula has to be considered as yielding a LFP-like signal.

### POPULATION SPIKE-TRIGGERED AVERAGE

We identified two cortical areas: a center area of 1mm radius aligned with the stimulus center, and a surround annulus area of 1 mm internal radius and 0.8 mm external radius. These two areas were alternatively used as “source” and “target of our analysis. We computed the LFP for the target location as described above; then we computed the population spike triggered average (STA), see supplementary figure 3. For each spike fired in the source location, we extracted an LFP chunk of 600 ms centered on the reference spike time, and we averaged across all extracted chunks. This STA was then analyzed to extract its features. We identified the presence of a negative peak as absolute STA minimum (“trough”) and characterized its lag, amplitude and duration. The lag was measured as (ms) difference between reference time and negative absolute minimum time. The amplitude was measured as the absolute minimum value (mV). And the duration was measured as the interval between the first two neighboring local maxima next to the absolute minimum (ms). We repeated this procedure for each stimulus size presentation.

### STATISTICAL SIGNIFICANCE ASSESSMENTS

The biological question we are asking is whether the LGN activities are changed by the cortical feedback. Given that we distinguished the experimental procedures into two configurations, control and variously inactivated feedback, we can formulate a null-hypothesis: “the LGN activities in the “full” and feedforward-only configurations are the same” and its corresponding alternative hypothesis. The experiments performed in the literature (that we replicated in-computo) identify two types of statistical tests depending on the kind of variable measured. For the ongoing activity and low-pass index of spatial frequency, we are dealing with a comparison of cells’ firing rates as measurement variables: therefore, one-way ANOVA (or Wilcoxon, for non normal distributions with skewness test not passed) was used. For all other experiments, where the firing rate was measured at different step changes of a stimulus value, the results were grouped into two categories: therefore, two-sample t-test (or Welch, for normal distributions with different standard deviations) was used. For the overlapping/non-overlapping surround suppression experiments, a non-parametric two-tailed test (Wilcoxon) was used, as in Jones et al. 2012. On the trial averaged firing rates, we also performed a statistical test for the null hypothesis “two related samples have identical average values” (as in Jones et al. 2012). In this case, the standard error of the mean was also computed.

### Software stack

In order to replicate as closely as possible the same experimental conditions as those in the literature, we need to have an experimental setup, not just a one-shot simulation. Therefore a whole infrastructure is needed, to prepare different types of stimuli, to operate selective inactivation of layers and units, to perform recordings and analysis.

We modeled the thalamo-cortical loop using Mozaik (Antolik and Davison 2013, see code at https://github.com/antolikjan/mozaik). Mozaik is an integrated workflow framework for large scale neural simulations, intended to relieve users from writing boilerplate code for projects involving complex heterogeneous neural network models, complex stimuli and experimental protocols and subsequent analysis and plotting. It is built on top of the following tools: *imagen* (for stimuli generation), *PyNN* 0.8.0 (for simulator independent neural network model definition, see http://neuralensemble.org/PyNN/), *NEST simulator* 2.1.0, *Neo* (for exchange and internal representation of data), *matplotlib* (for plotting). Additional analysis code was written using the *scipy* library (https://www.scipy.org/). All the code is available online (see https://github.com/dguarino/T2). Experimental data were digitized by scraping the original published material using WebPlotDigitizer 3.11 (http://arohatgi.info/WebPlotDigitizer).

## Acknowledgments

We would like to thank Damien Depannemaecker, Karolína Korvasová and Alain Destexhe, for stimulating discussions and critically reading the manuscript.

This work was primarily done at the UNIC (Unit of Neuroscience, Information and Complexity, UPR CNRS 3293). The work was funded by grants to Y.F. and A. D. from the Agence Nationale de la Recherche (ANR-Horizontal-V1 (ANR-17-CE37-0006), the Paris-Saclay IDEX (NeuroSaclay and i-Code) and the FET-Proactive Grant Agreement No 269921 (BrainScales). D.G. received continuous funding from the EC Human Brain Project (HBP, grant agreement, n. 604102). The work was also funded by grant to J.A. from Charles University in Prague (PRIMUS/20/MED/006) and joined grant to J.A. and D.G. (CZ MSMT project No. 8J20FR006).

---

**Figure S1:**
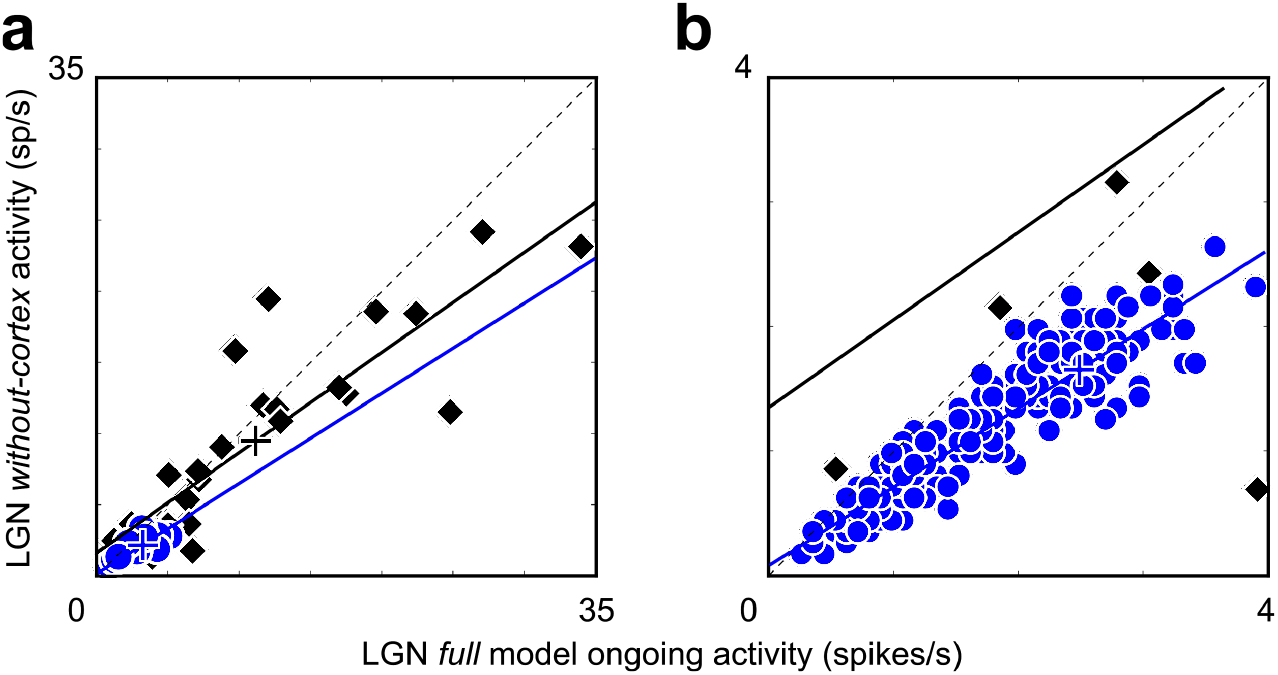
Cortical feedback reduces ongoing activity in the model LGN. (**a**) Comparison of LGN ongoing activity in the “without-cortex” (vertical axis) vs “full” (horizontal axis) models. Model data (blue circles) follow a decrease slope similar to the experimental data from Waleszczyk et al. 2005 (black diamonds). This decrease is found significant in the simulation case (Kruskal-Wallis p<0.001). Linear regressions are plotted for the data in black, and for the model in blue. The dotted line represents identity (no change between configurations). (**b**) Zoomed portion of the same data as in (**a**) to better appreciate model data points distribution and their deviation from the identity line.

**Figure S2:**
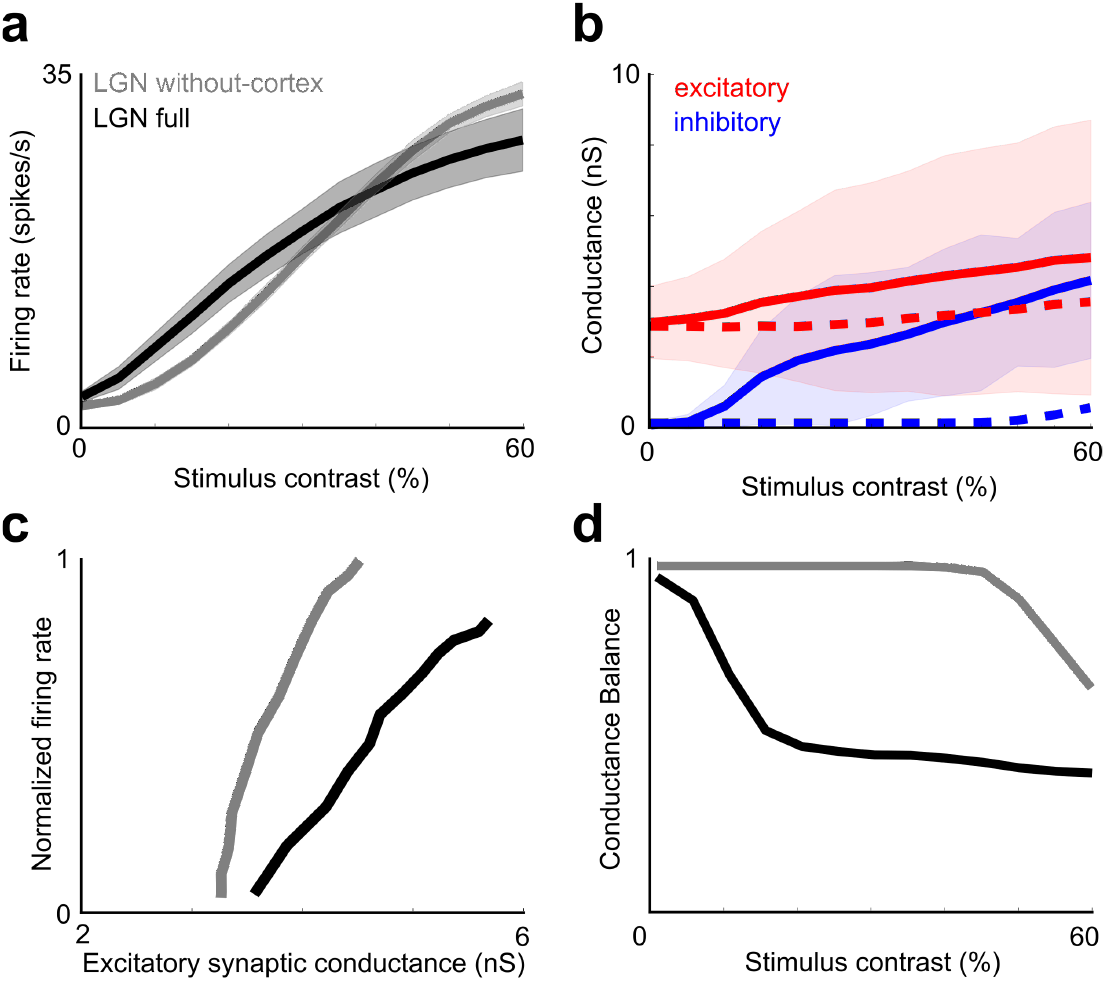
Cortical feedback changes contrast sensitivity of LGN cells. (**a**) Contrast tuning curves based on trial averaged firing rates for the On-center LGN population (Off-center follows the same trend which has not been overlaid for clarity). The comparison between “full” (black mean lines and shaded SEM) and “without-cortex” (grey mean line and shaded SEM) configurations show that cortical feedback increases thalamic firing rate for low-contrast stimuli (c<0.4) and reduces it for high-contrast stimuli (c>0.4) (grey mean line and shaded SEM). (**b**) Input-output ratio for LGN On-center cells computed using trial-averaged conductances and trial-averaged firing rates. In the “without-cortex” model (grey), the dynamic range is significantly reduced compared to the “full” model (black). (**c**) Trial-averaged excitatory (red) and inhibitory (blue) conductance tuning curves for On-center LGN cells (Off-center cells shared the same properties) in the “full” (solid) and “without-cortex” (dashed) models. Both evoked excitatory and inhibitory conductances increase with the stimulus contrast and with the cortical feedback, but the increase slopes differ, being faster for inhibitory input than for excitatory drive. (**d**) This results in a striking difference in the contrast dependency for the balance between excitation and inhibition, where inhibitory damping prevails in the “full model” as soon as the contrast increases above 20%.

**Figure S3:**
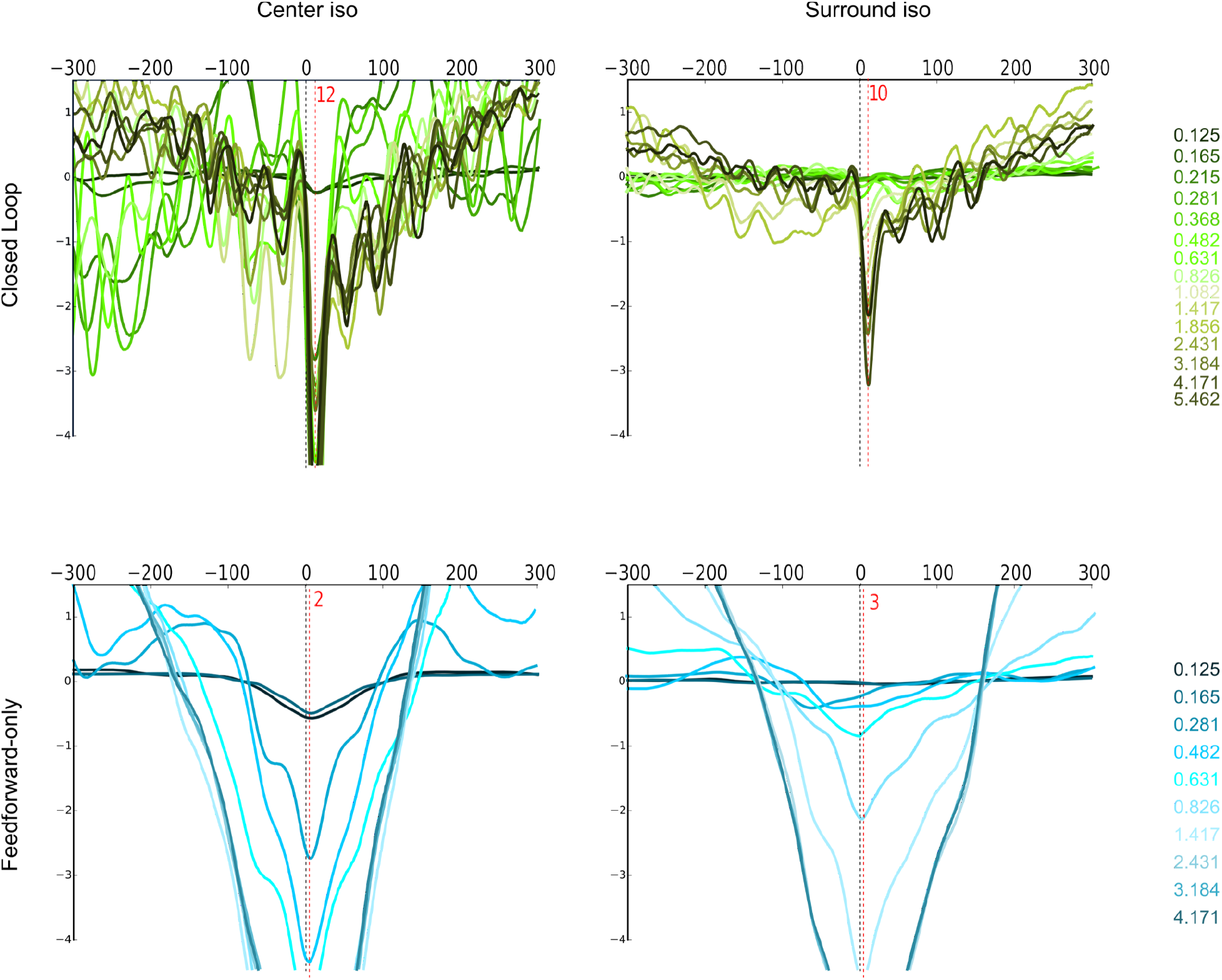
Overlayed Population Spike-Triggered Averages for all stimulus sizes in the center and surround. (**Top row, Left**) PSTA of synthetic LFP recorded in the center iso group triggered by spikes emitted in the surround, in the full model (closed loop) configuration, for different sizes (color code on the left corresponding to the radius). The PSTA amplitude is tuned for stimulus size (see main text figure 8a). The PSTA width is constant across sizes. And the PSTA lag is reliably following the triggering spikes (∼12ms). (**Top row, Right**) PSTA of sLFP recorded in the surround iso group triggered by spikes emitted in the center, in the full model (closed loop) configuration, for different sizes (color code on the left corresponding to the radius). The PSTA amplitude is tuned for stimulus size (see main text figure 8b). The PSTA width is constant across sizes. And the PSTA lag is reliably following the triggering spikes (∼10ms). (**Bottom row, Left**) PSTA of sLFP recorded in the center iso group triggered by spikes emitted in the surround, in the feedforward-only model configuration, for different sizes (color code on the left corresponding to the radius). (**Bottom row, Right**) PSTA of sLFP recorded in the surround iso group triggered by spikes emitted in the center, in the feedforward-only model configuration, for different sizes (color code on the left corresponding to the radius). For both lower panels, the PSTA amplitude is not tuned for stimulus size (see main text figure 8c-d). The PSTA width is monotonically growing across sizes. And the PSTA lag is smaller (∼2ms) compared to the full model.

**Figure S4:**
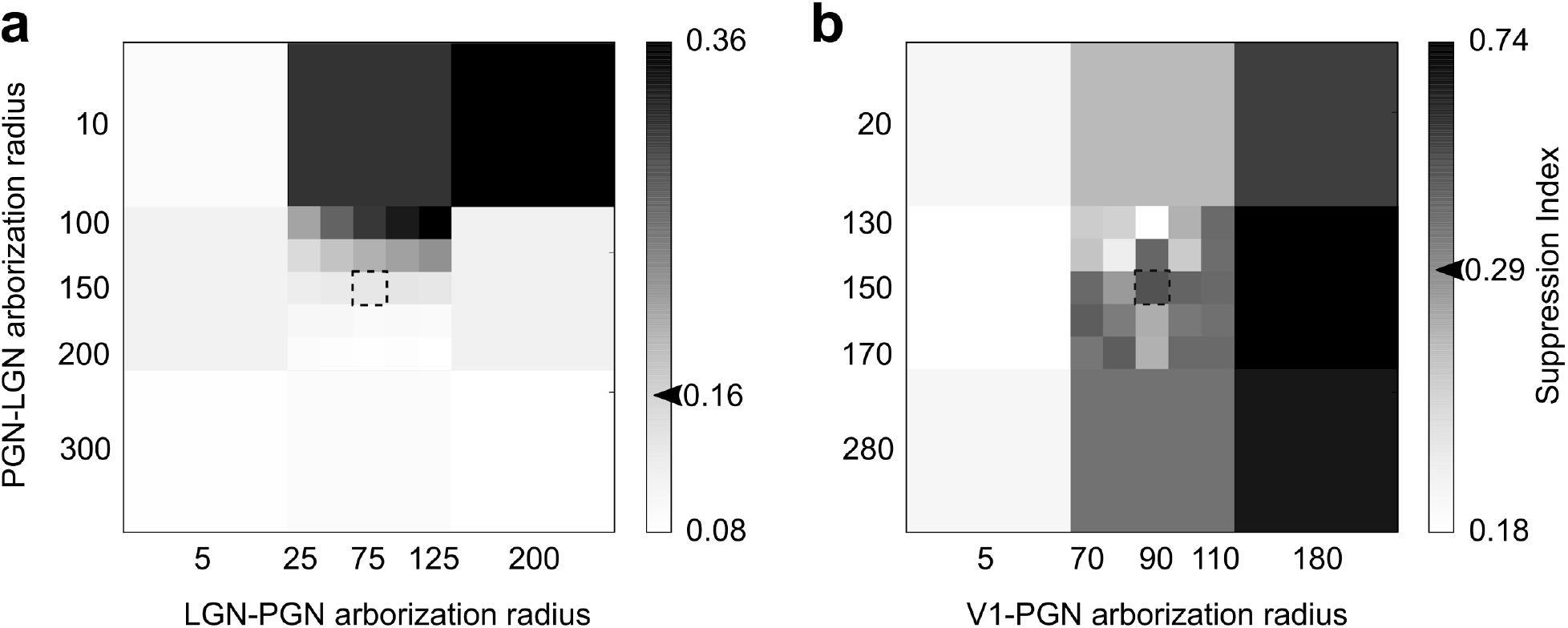
Parameter searches to assess stability of intra-thalamic and corticothalamic arborizations. (**a**) Suppression index (color code on the bar) resulting from changing the arborisation extent (σ) of LGN→PGN and PGN→LGN for the without-cortex configurations. (**b**) Suppression index resulting from changing the arborisation extent (σ) of V1→PGN for the full and without-cortex configurations. In both figures, the dashed square in the middle is the chosen parameter combination, and the arrowheads on the color bars are reference values from the literature (**a**: Jones and Sillito 1987; **b**: Jones et al. 2012).

